# Haplotype-resolved genome assembly of barberry reveals structural divergence and allele-specific expression during infection by wheat stripe rust pathogen

**DOI:** 10.64898/2026.09.01.748723

**Authors:** Jierong Wang, Yiwen Xu, Yumeng Duan, Zhensheng Kang, Jing Zhao

## Abstract

Barberry is an ecologically and medicinally important perennial shrub and an alternate host of *Puccinia striiformis* f. sp. *tritici* (*Pst*), the causal agent of wheat stripe rust. However, its genome architecture and molecular responses to *Pst* infection remain poorly understood. Here, we generated a haplotype-resolved, chromosome-level genome assembly of *Berberis aggregata* using PacBio HiFi and Hi-C sequencing. The two haplotypes, Ba5A and Ba5B, were assembled into 14 pseudochromosomes each, with genome sizes of approximately 1.13 and 1.16 Gb, respectively, and showed high continuity and completeness. Comparative analyses revealed extensive divergence between the haplotypes, including widespread structural variation, substantial hemizygosity, and TE-rich, gene-poor non-alignable regions. Haplotype-specific hemizygous genes also exhibited distinct functional biases, with Ba5A enriched in immune-related processes and Ba5B associated with transport and cell-envelope functions. Dual RNA-seq across infection stages revealed stage-dependent host transcriptional reprogramming, characterized by early pathogen perception followed by stronger defense activation and photosynthesis repression, whereas *Pst* displayed a coordinated colonization program consistent with its biotrophic lifestyle. Allele-specific expression analysis showed that ∼25–28% of genes exhibited significant haplotype-biased expression, and a subset transitioned from unbiased expression at 0 dpi to Ba5A- or Ba5B-biased expression during infection. Integration with differential expression analysis further identified infection-responsive induced-biased genes associated with defense-related processes, including responses to fungal pathogens. These findings show that haplotype divergence and dynamic allele-specific regulation jointly contribute to transcriptional responses during *Pst* infection, providing new insights into the genomic basis of barberry–rust pathogen interactions.

## 1. Introduction

Barberry (*Berberis spp.*) is a perennial woody shrub widely distributed in temperate and subtropical regions, and is taxonomically well represented in the Berberidaceae family (Rounsaville and Ranney 2010). It is valued both as an ecological landscape plant and as a medicinal resource due to its rich accumulation of bioactive alkaloids, such as berberine, which have been widely used in traditional medicine (Sobhani *et al*. 2021; Bathaei *et al*. 2025). In addition to its ecological and pharmacological importance, barberry serves as alternate hosts for several cereal rust fungi, most notably *Puccinia striiformis* f. sp. *tritici* (*Pst*) and *Puccinia graminis* f. sp. *tritici* (*Pgt*), the causal agents of wheat stripe rust and wheat stem rust, respectively (Zhao *et al*. 2016). These diseases are among the most destructive threats to global wheat production, with annual economic losses caused by wheat rust pathogens estimated at US$4.3–5.0 billion (Figueroa *et al*. 2018; Singh *et al*. 2025). Completion of the sexual stage of *Pst* and *Pgt* on susceptible barberry hosts facilitates genetic recombination and may promote the emergence of novel genotypes and virulent races, making barberry and cereal rust fungus interactions an important component of rust disease epidemiology and pathogen evolution (Duplessis *et al*. 2021; Zhao and Kang 2023). Despite the ecological, agricultural, and medicinal importance of barberry, its genomic characteristics and the molecular mechanisms underlying its responses to rust fungal infection remain largely unexplored.

Plant responses to pathogen infection involve extensive and dynamic transcriptional reprogramming, including the activation of immune signaling, metabolic reconfiguration, and coordinated adjustment of growth-related processes (Huot *et al*. 2014; Jian *et al*. 2024). In diploid and highly heterozygous species, most gene loci are represented by two alleles located on homologous chromosomes, and sequence or regulatory differences between these alleles may result in unequal transcriptional activity. Allele-specific expression (ASE), defined as the preferential expression of one allele over the other within the same genetic and cellular environment, commonly arises from cis-regulatory sequence variation or allele-associated epigenetic differences (Pastinen 2010). ASE provides a direct link between genetic variation and transcriptional output and has been associated with phenotypic diversity, environmental adaptation, and stress-responsive plasticity (Albert and Kruglyak 2015; Moyerbrailean *et al*. 2016; Albert *et al*. 2018). For example, insertion of a transposable element into one allele of the flowering regulator *TFL* in strawberry caused allele-specific silencing and contributed to the transition from seasonal to perpetual flowering (Iwata *et al*. 2012). In apple, approximately 19% of expressed genes showed ASE, and many of these genes, including those encoding ACC oxidase and RIN-like MADS-box transcription factors, were associated with fruit development and quality traits (Sun *et al*. 2020). Similarly, haplotype-resolved transcriptome analysis in the tea plant identified numerous genes exhibiting salt- or drought-responsive ASE, including genes associated with ubiquitination, phenylpropanoid metabolism, and stress signaling (Zhang *et al*. 2023). Together, these findings indicate that ASE can provide a flexible regulatory mechanism through which heterozygous plants adjust gene expression in response to developmental and environmental cues. However, most studies of plant–pathogen interactions have focused primarily on gene-level expression changes, and the contribution of allele-specific regulatory variation to infection-responsive transcriptional reprogramming remains poorly understood.

With advances in sequencing technologies and bioinformatic tools, increasingly complex plant genomes can now be assembled at haplotype-resolved and even telomere-to-telomere (T2T) levels (Li and Durbin 2024). Such high-quality assemblies enable the resolution of chromosome-scale genome organization, structural divergence between haplotypes, and haplotype-specific gene content, thereby providing new opportunities to understand biological differentiation and plant–microbe interactions (Cheng *et al*. 2021; Shi *et al*. 2024). Haplotype-resolved genomes also provide an essential foundation for accurate ASE analyses by allowing reads to be assigned to their corresponding alleles and enabling systematic dissection of regulatory divergence between haplotypes (Cheng *et al*. 2021; Han *et al*. 2024; Shi *et al*. 2024). Previous studies have shown that allelic expression imbalance can be highly dynamic and condition-dependent, particularly in response to environmental stress and immune activation (Wu *et al*. 2026). However, genomic resources for barberry remain limited, and previously published barberry genome assemblies have generally lacked sufficient haplotype resolution to distinguish homologous chromosomes and characterize haplotype-specific gene content accurately (Bartaula *et al*. 2019; Ruhsam 2024). This limitation hinders the reliable assignment of transcriptomic reads to individual alleles and constrains genome-wide investigation of ASE during rust pathogen infection. Consequently, how haplotype-resolved genomic architecture shapes infection-responsive transcriptional reprogramming in barberry remains largely unknown.

Here, we generated a high-quality haplotype-resolved genome assembly of *Berberis aggregata* Schneid. and performed dual RNA sequencing across multiple *Pst* infection stages. We systematically characterized genome-wide structural variation, transcriptional dynamics of both host and pathogen, and ASE patterns during pathogen infection. By integrating differential expression and haplotype-biased expression analyses, we identified genes showing infection-induced shifts from unbiased to haplotype-biased expression, and further explored their functional roles in host–pathogen interactions. Our results provide an integrated genomic and transcriptomic framework for understanding the interaction between barberry and stripe rust fungus.

## 2. Materials and methods

### 2.1. DNA extraction, sequencing, and genome size estimation of *B. aggregata*

Genomic DNA was extracted from young leaves of *B. aggregata* with the previously described method (Schwessinger and Rathjen 2017). For PacBio HiFi sequencing, a SMRT bell library was constructed and sequenced on PacBio Sequel II system. Meanwhile, a DNA library with 350-bp fragment sizes was constructed and sequenced using Illumina Novaseq PE150 platform. The Hi-C library was constructed using the 4-cutter restriction enzyme *DpnII* and sequenced on the Illumina Novaseq PE150 platform. RNA was extracted separately from young *B. aggregata* leaves using the QIAGEN (Doncaster, Australia) Plant RNeasy kit as described previously (Zhao *et al*. 2021a). Equal amounts of the three RNA samples were mixed for mRNA sequencing by Illumina Novaseq sequencing.

Before assembly, the genome size and heterozygosity were estimated with Illumina short DNA reads. Jellyfish v2.3.0 (Marçais and Kingsford 2011) was used to calculate the frequency distribution of the depth of clean data with 29-mer. Then the results were imported to GenomeScope v2.0 (Vurture *et al*. 2017) to estimate the basic features of the genome with 29-mer.

### 2.2. Genome assembly and assessment of *B. aggregata*

PacBio HiFi reads and Hi-C short reads were used as a combined input for the genome assembly using Hifiasm v0.19.0 (Cheng *et al*. 2021) in Hi-C mode with the default parameters. Hi-C paired-end clean reads were aligned to the assembly using Juicer v1.6.2 (Durand *et al*. 2016) with the BWA algorithm to get the interaction matrix. The 3d-DNA v180922 pipeline (Dudchenko *et al*. 2017) was applied to reorder and scaffold the contigs. The position of the contigs was also manually adjusted based on Hi-C heatmaps visualized with JuicerBox v1.9.8 (Robinson *et al*. 2018).

Blastn searches against the NCBI nr/nt database were used to check potential contamination and none of the contigs were significant hits to noneukaryotic sequences, chloroplast sequences, mitochondrial sequences or plant rRNA with E-value set as 1e-10. The obtained contigs were parsed by Purge Haplotigs v1.1.1 (Roach *et al*. 2018) and Redundans (Pryszcz and Gabaldón 2016) to get rid of the redundancies. The TGS-GapCloser v1.2.1 (Xu *et al*. 2020) was utilized to fill gaps based on the haplotype-resolved chromosome-level genome, using the PacBio HiFi sequencing data. The telomere identification tool (https://github.com/tolkit/telomeric-identifier) was used to search for the normalized unified sequence TTTAGGG/CCCTAAA within 50 kb of each terminal chromosome sequence.

The quality of genome assembly was evaluated with multiple methods. First, the accuracy of Hi-C based chromosome construction was evaluated by chromatin contact matrix with HiC-Pro v3.0.0 (Servant *et al*. 2015), and contact maps were plotted with hicPlotMatrix of HiCExplorer v3.7.2 (Ramírez *et al*. 2018). Second, the BUSCO analysis with the embryophyta_odb10 database (genome mode) was performed to assess genome completeness using BUSCO v5.4.7 (Simão *et al*. 2015). Third, Illumina short reads and HiFi long reads from AZ2 were mapped to the assembly using BWA-MEM (Li and Durbin 2009) and minimap2 v2.24 (Li 2018), then QualiMap v2.2 (Okonechnikov *et al*. 2016) was used to evaluate the mapping quality. Forth, the consensus quality value (QV) and completeness of the genome was evaluated using Merqury v1.3 (Rhie *et al*. 2020) with meryl v1.3 (under 19-mer) count.

### 2.3. Gene prediction and functional annotation of *B. aggregata*

EDTA pipeline v2.1.0 (Ou *et al*. 2019) was used to annotate the repetitive sequences in the *B. aggregata* genome with parameters “--step all --sensitive 1 --anno 1 --evaluate 1”. Protein-coding genes were predicted by integrating ab initio prediction and transcriptome-based prediction. RNA-seq data sets (Zhao *et al*. 2021b) were aligned to the soft-masked assembly using Hisat2 v2.2.1 (Kim *et al*. 2015). StringTie v2.2.1 (Pertea *et al*. 2015) was used to reconstruct the transcript, and TransDecoder v5.7.1 (https://github.com/TransDecoder/TransDecoder) was used to predict the gene models. In addition, Trinity v2.15.1 (Grabherr *et al*. 2011) was used to assemble the transcriptome, and then the assembled transcripts were used for gene structure prediction with PASA v2.5.3 (Haas 2003). Ab initio gene prediction was carried out using BRAKER v2.1.6 (Brůna *et al*. 2021), incorporating proteins from embryophyta_odb10 database and the mapping data of the RNA-seq reads. All the above gene models were combined using EvidenceModeler v.1.1.1 (Haas *et al*. 2008) with default weight settings. The completeness of the gene prediction was evaluated using BUSCO v5.4.7 (Simão *et al*. 2015) with the embryophyta_odb10 database (protein mode).

Functional annotation of the protein-coding genes was performed in multiple databases, including Kyoto Encyclopedia of Genes and Genomes (KEGG), and Cluster of Orthologous Groups of proteins (KOG/COG/eggNOG) based on eggNOG-mapper v2.1.12 (Huerta-Cepas *et al*. 2019; Cantalapiedra *et al*. 2021); UniProtKB/SwissProt and NCBI non-redundant protein (Nr) by using diamond v2.1.8.162 (Buchfink *et al*. 2015) with an e-value threshold of 1e-5; and Pfam by applying InterProScan v5.74-105.0 (Jones *et al*. 2014). The Gene Ontology (GO) annotation was created by combining the results of eggNOG-mapper v2.1.12 (Huerta-Cepas *et al*. 2019; Cantalapiedra *et al*. 2021) and InterProScan v5.74-105.0 (Jones *et al*. 2014).

### 2.4. Variation detection between two haplotypes of *B. aggregata*

To investigate the differences between the two haplotypes, the command nucmer in MUMmer v4.0 (Marçais *et al*. 2018) with the parameters ‘--maxmatch -c 100 -l 500’ was used for whole-genome alignments, and the alignment results were filtered using the command delta-filter with the parameters ‘-m -i 90 -l 100’. After format conversion with the command show-coords, SyRI v1.6.3 (Goel *et al*. 2019) with the default parameters was used to detect syntenic regions and structural variations. Plotsr v1.1.1 (Goel and Schneeberger 2022) was used to visualize the variations.

Protein sequences from all annotated genes were analyzed using Proteinortho v6.3.5 (Lechner *et al*. 2011) with the -synteny flag to detect homologs between haplotypes. Genes lacking a hit on the alternative haplotype were considered hemizygous candidates. These were further filtered via reciprocal BLASTp to ensure the absence of alleles. Candidates with a high-quality hit that had >70% identity and >70% query and subject coverage were omitted.

### 2.5. Dual RNA-seq read mapping and expression quantification

RNA-seq reads were mapped to a combined reference consisting of the Ba5A genome of *B. aggregata* and the AZ2B genome of *Pst* isolate AZ2 (Wang *et al*. 2024) using Hisat2 v2.2.1 (Kim *et al*. 2015). Host- and pathogen-assigned read proportions were calculated from the combined alignments. Gene-level expression was quantified as raw counts using StringTie v2.2.1 (Pertea *et al*. 2015), and normalized expression values (FPKM) were calculated for visualization. Principal component analysis (PCA) was performed separately for host and pathogen transcriptomes using normalized expression matrices to evaluate sample clustering and stage-specific transcriptional shifts.

### 2.6. Differential expression analysis and functional enrichment analyses

Differential expression analyses for the host were performed by comparing each infected time point (3, 4, 7 and 11 dpi) with the uninfected control (0 dpi). Differential expression analyses for the pathogen were performed across successive infection-stage transitions (4 vs 3 dpi, 7 vs 4 dpi and 11 vs 7 dpi). Raw count matrices were filtered to remove lowly expressed genes, and differential expression was tested using DESeq2 (Love *et al*. 2014). Genes with an adjusted-*P*-value <0.05 and |log2FoldChange| ≥1 were considered differentially expressed.

GO biological process and KEGG pathway enrichment analyses were performed separately for upregulated and downregulated gene sets using clusterProfiler v4.4.4 (Wu *et al*. 2021). For barberry, enriched GO terms were grouped into broader biological modules, including defense, hormone signaling, perception/signaling, transport, cell wall/structure and metabolism, to facilitate interpretation. For *Pst*, enriched GO terms were grouped into modules related to carbohydrate metabolism and nutrient utilization, homeostasis and adaptation, growth/development, and translation/RNA processing/protein biogenesis. Enrichment significance was evaluated using adjusted-*P*-values.

### 2.7. Allelic gene expression analysis

RNA-seq reads were mapped to a combined reference comprising the Ba5A and Ba5B genomes of *B. aggregata* using Hisat2 v2.2.1 (Kim *et al*. 2015), and only uniquely mapped reads were retained. ASE was analyzed for one-to-one Ba5A/B alleles using DESeq2 (Love *et al*. 2014). At each time point, alleles with a combined Ba5A and Ba5B count of at least six reads in all three biological replicates were retained, whereas the remaining alleles were classified as low-expression. Alleles with an adjusted-*P*-value <0.05 and |log2FoldChange| ≥2 were classified as showing ASE. These ASE genes were further classified as Ba5A-biased or Ba5B-biased when the log2FoldChange was ≥2 or ≤-2, respectively. Alleles that passed the expression filter but did not meet both criteria were operationally classified as unbiased.

## 3. Results

### 3.1. A haplotype-resolved chromosome-level genome assembly of *B. aggregata*

We performed the genome sequencing and de novo assembly of *B. aggregata* by combining PacBio HiFi, Illumina, and Hi-C technologies. Initially, the *B. aggregata* genome was estimated using 63.45 Gb Illumina sequencing data by *K*-mer. The estimated genome size of *B. aggregata* was 974.04 Mb, and the heterozygosity rate was 0.95% (Fig. 1A). The PacBio HiFi platform was then used to generate 97.35 Gb data with an average read length of 18.11 Kb and a sequencing depth of approximately 115× (Table S1). In addition, 100.26 Gb of Hi-C data (∼118×genome coverage) was generated using the PE150 sequencing. Preliminary haplotype-resolved assemblies were generated using HiFi reads and the Hi-C mode of Hifiasm, followed by chromosome-scale scaffolding with Hi-C data. Each haplotype was successfully anchored into 14 pseudo-chromosomes (Fig. 1B-C). After gap filling, the two haplotype assemblies, designated Ba5A and Ba5B, had total sizes of 1.13 Gb and 1.16 Gb, respectively, with scaffold N50 values of 82.45 Mb and 82.59 Mb (Table 1). Only 15 gaps were detected in Ba5A and 25 in Ba5B, indicating a high level of assembly continuity. Telomeric repeat searches (TTTAGGG/CCCTAAA) identified 55 telomeres across the two haplotypes, with telomeric sequences present at both ends of 27 chromosomes and at one end of one chromosome (Fig. 1D). Together, these results indicate that a highly continuous haplotype-resolved chromosome-level assembly of *B. aggregata* was successfully generated, providing a robust genomic reference for subsequent analyses.

**Fig. 1.**
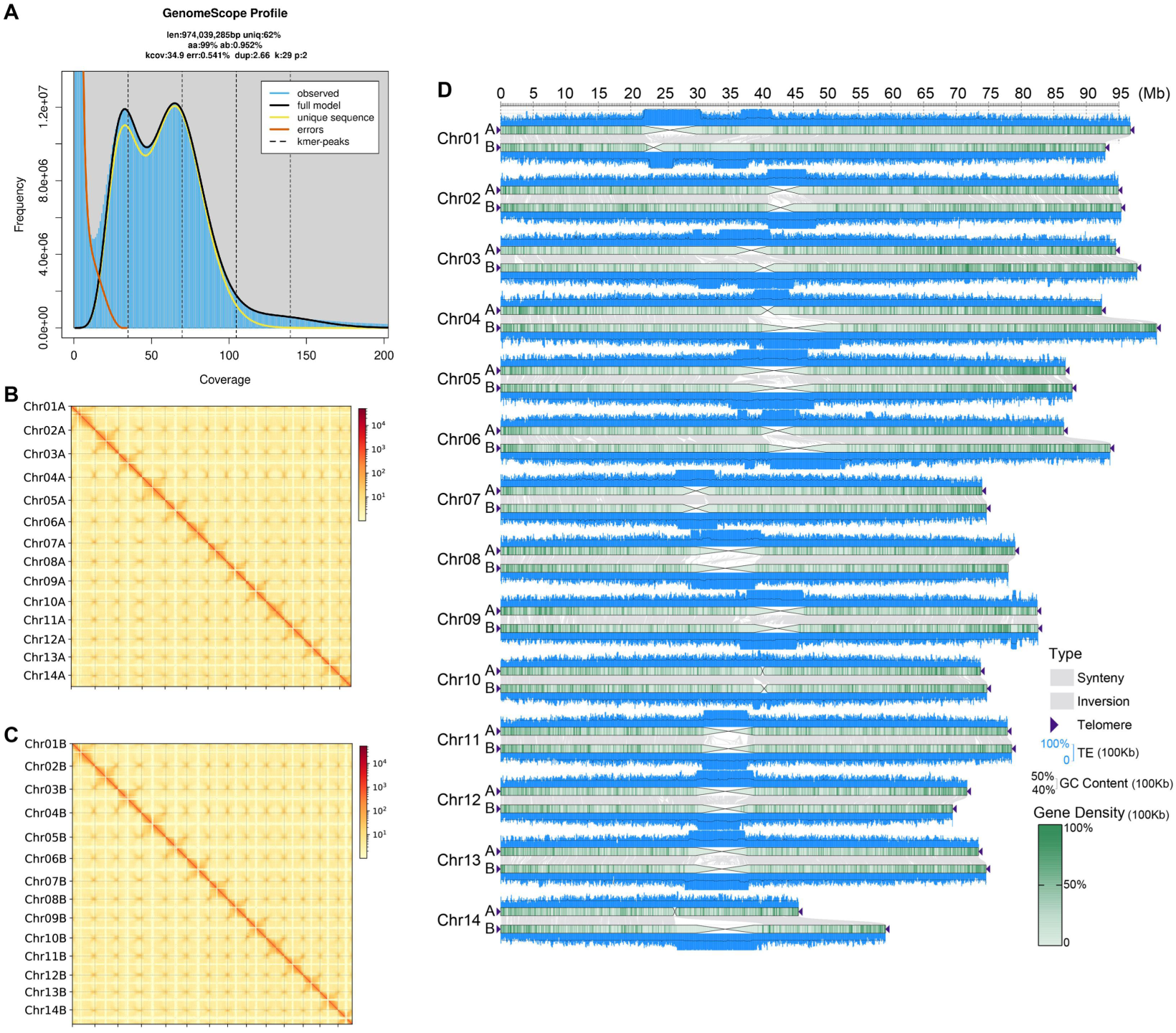
Haplotype-resolved chromosome-level genome assembly of *Berberis aggregata*. A, GenomeScope profile based on Illumina *k*-mer analysis, showing the estimated genome size and heterozygosity of *B. aggregata*. B and C, Hi-C contact maps of the two phased haplotypes, Ba5A and Ba5B, respectively, showing strong intrachromosomal interaction signals and successful anchoring of each haplotype into 14 pseudochromosomes. D, Chromosome-scale overview of the two haplotype assemblies, illustrating the organization of the 14 pseudochromosomes in Ba5A and Ba5B and the overall correspondence between homologous chromosomes. Together, these results support a high-quality haplotype-resolved chromosome-level assembly of *B. aggregata*.

**Table 1.**
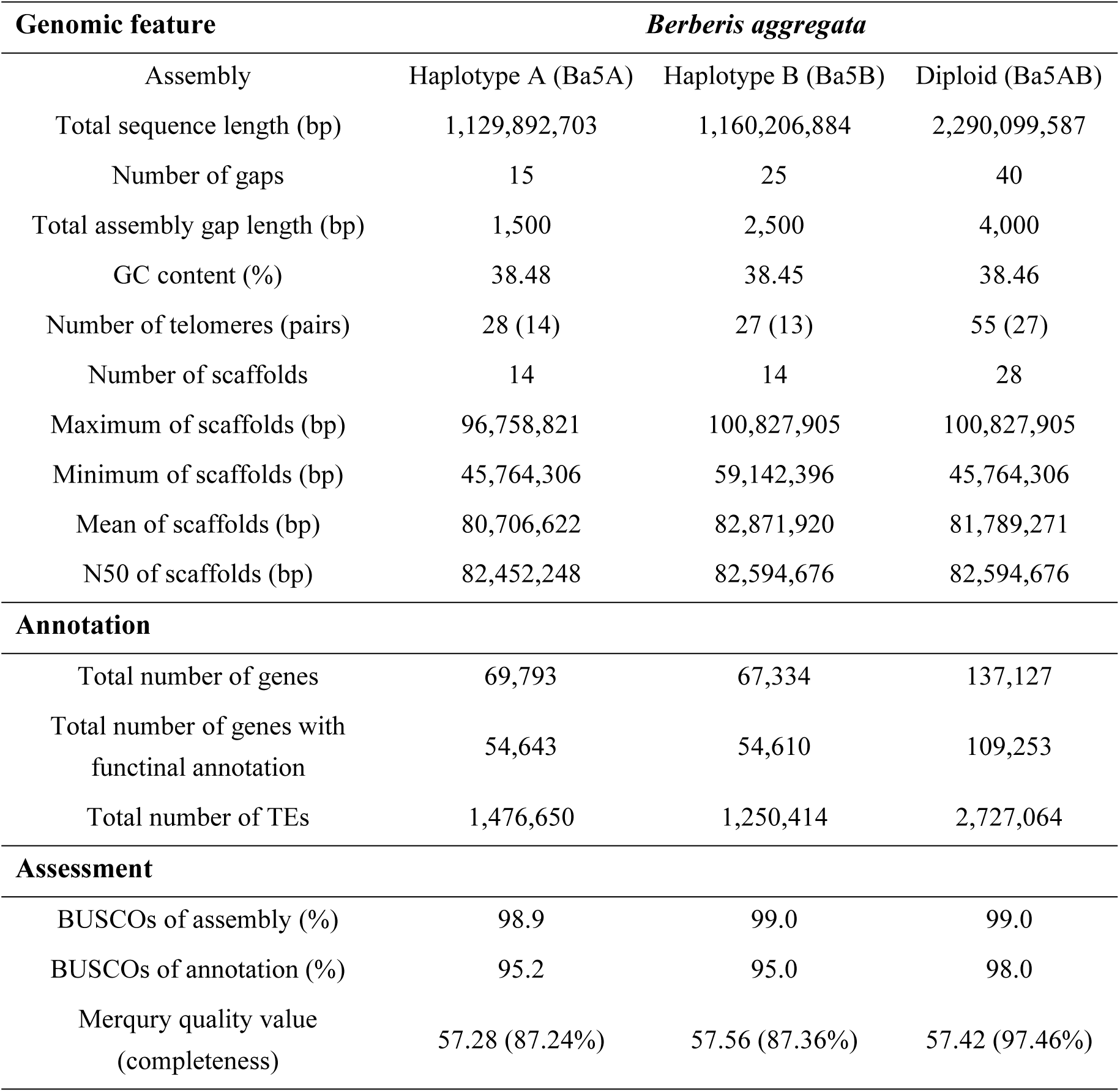
Characteristics of *Berberis aggregata* genome assembly.

Multiple complementary approaches were applied to assess the quality of the *B. aggregata* genome assembly. Illumina short reads showed a mapping rate of 99.38% and covered 99.94% of the assembled genome (Table S2). PacBio HiFi reads exhibited an even higher mapping rate of 99.96%, with 100% genome coverage. Merqury analysis further demonstrated the high consensus accuracy of the assembly, yielding a Quality Value (QV) of 57.42 (Table S3). The haplotype-specific QV scores were 57.28 for Ba5A and 57.56 for Ba5B, corresponding to an estimated base accuracy of 99.999%. Genome completeness was evaluated using BUSCO with 1,614 conserved embryophyte orthologs. Overall, 99.0% of BUSCO genes were completely recovered in the combined assembly (Table S4). The Ba5A and Ba5B haplotypes individually achieved BUSCO completeness scores of 98.9% and 99.0%, respectively. Together, these metrics demonstrate the high accuracy, completeness, and effective haplotype phasing of the *B. aggregata* genome assembly.

### 3.2. Genome annotation and repeat composition

A comprehensive annotation of repetitive elements identified 1,476,650 and 1,250,414 transposable elements (TEs) and repeats in Ba5A and Ba5B, accounting for 63.03% and 64.21% of the respective assemblies (Table S5). Long terminal repeat (LTR) retrotransposons were the most abundant TE class, comprising 30.47% of Ba5A and 28.70% of Ba5B. Among these, Gypsy elements predominated, representing 16.20% and 15.93% of Ba5A and Ba5B, respectively. Terminal inverted repeat (TIR) elements constituted another major TE class, accounting for 23.28% of Ba5A and 24.92% of Ba5B.

A total of 137,127 protein-coding genes were predicted, including 69,793 genes in Ba5A and 67,334 genes in Ba5B. BUSCO assessment of the gene models indicated a completeness of 98.0% for the annotated gene set (Table S6), with complete BUSCO scores of 95.2% and 95.0% for Ba5A and Ba5B, respectively. Owing to its slightly higher annotation completeness and continuity, Ba5A was used as the reference haplotype for subsequent comparative genomic analyses and gene expression quantification. Functional annotation against multiple protein databases assigned putative functions to 54,643 genes (78.29%) in Ba5A and 54,610 genes (81.10%) in Ba5B (Fig.S1A-B and Table S7), facilitating downstream analyses of haplotype-specific gene content and host–pathogen transcriptomic responses.

### 3.3. Extensive inter-haplotype structural divergence and hemizygosity

To characterize inter-haplotype genomic variation, we aligned the Ba5B assembly against Ba5A. Approximately 80% of the two haplotypes exhibited collinear synteny (Fig. 2A), with 901.38 Mb (79.78%) of Ba5A and 900.06 Mb (77.58%) of Ba5B forming highly continuous syntenic blocks. We identified a total of 9,575 and 7,724 chromosomal structural variations (SVs) in Ba5A and Ba5B, respectively, including 111 inversions, 2,758 translocations, 8,991 indels (≥ 50 bp), and extensive duplications (6,706 in Ba5A and 4,855 in Ba5B) (Fig. 2B, Table S8). These SVs collectively accounted for approximately 9.57% of the Ba5A genome and 7.76% of the Ba5B genome. In addition, 13.20% of Ba5A and 15.87% of Ba5B sequences were non-alignable between haplotypes, indicating a substantial proportion of hemizygous regions. Most syntenic blocks ranged from 10 to 1,000 kb in length, while different SV types exhibited length distributions predominantly between 1 and 100 kb (Fig. 2B).

**Fig. 2.**
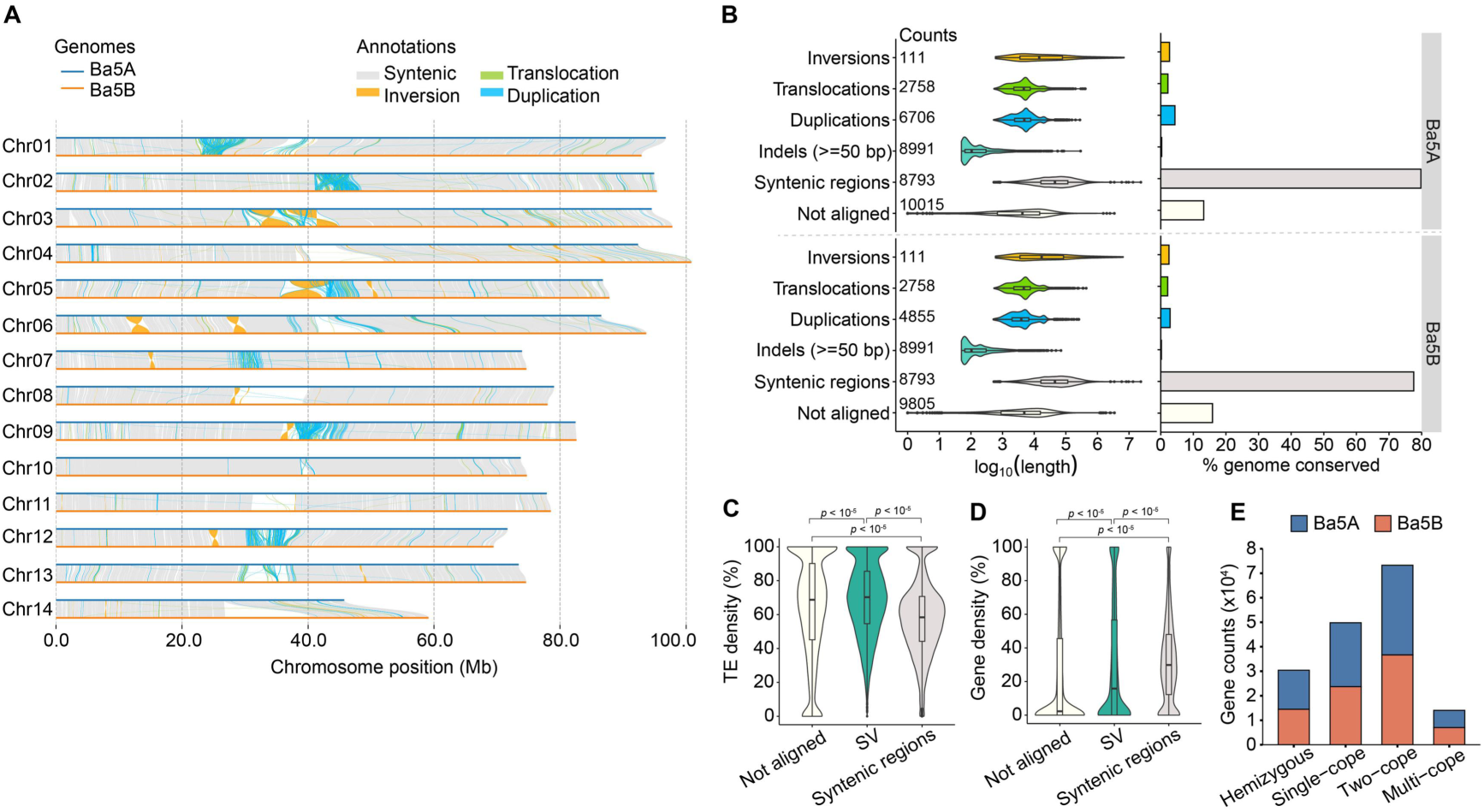
Extensive inter-haplotype structural divergence and hemizygosity between Ba5A and Ba5B. A, Genome-wide pairwise comparison of the two haplotypes. Homologous chromosomes of Ba5A and Ba5B are shown as horizontal bars, and aligned regions are classified as collinear syntenic blocks, inversions, translocations and duplications. B, Summary of inter-haplotype structural variation, showing the counts and length distributions of different categories of aligned regions and structural variants in Ba5A and Ba5B. C and D, Distributions of TE density and gene density in syntenic, SV and non-alignable regions. Compared with syntenic regions, structurally variable and non-alignable regions are enriched in transposable elements and depleted in genes. E, Classification of genes according to orthology relationships between the two haplotypes, including hemizygous, single-copy, two-copy and multi-copy genes in Ba5A and Ba5B.

To further characterize the sequence composition of inter-haplotype variation, we compared TE and gene densities among syntenic, SV, and hemizygous regions (Fig. 2C and D). TE density was significantly higher in SV and especially in non-alignable regions than in syntenic regions, whereas gene density showed the opposite trend, being most enriched in syntenic blocks and markedly reduced in SV and hemizygous regions (Wilcoxon test, *p* < 2.2 × 10^−16^). These patterns indicate that inter-haplotype divergence in *B. aggregata* is closely associated with TE accumulation and local gene depletion, with hemizygous regions representing repeat-rich and gene-poor compartments that likely contribute to structural and functional differentiation between haplotypes.

To assess the impact of haplotype divergence on gene content, orthology analysis using Proteinortho (Lechner *et al*. 2011) identified 87,330 genes with homologs in the alternative haplotype. These included 73,282 duplicated genes (36,641 one-to-one pairs) and 14,048 multi-copy genes (Fig. 2E). In addition, 49,797 single-copy genes were detected (26,107 in Ba5A and 23,690 in Ba5B). Reciprocal BLASTP analysis further identified 15,980 (22.90%) and 14,456 (21.47%) hemizygous genes unique to Ba5A and Ba5B, respectively, consistent with the extensive hemizygosity observed at the genomic level. GO enrichment analysis further revealed clear haplotype-specific functional biases, with Ba5A-unique genes enriched in immune-related processes and Ba5B-unique genes preferentially associated with transport- and cell envelope-related functions, including ion transport and suberin biosynthesis, indicating functional partitioning between haplotypes that may contribute to host–pathogen interactions (Fig. S2, Table S9). Together, these results suggest that the two haplotypes are not only structurally differentiated, but also functionally partitioned, with TE-associated hemizygosity likely playing an important role in shaping haplotype-specific gene content and biological specialization.

### 3.4. Stage-specific host–pathogen transcriptional dynamics during *Pst* infection of barberry

To characterize the transcriptional dynamics of both host and pathogen during *Pst* infection, RNA-seq was performed using barberry samples collected at 0, 3, 4, 7, and 11 days post-inoculation (dpi). Across all libraries, 159–232 million paired-end reads were generated per sample, with 79.6%–85.9% uniquely mapped to the reference genomes (Table S10). Mapping depth was broadly consistent among biological replicates, indicating high sequencing quality and comparable transcriptome coverage across infection stages. Dual RNA-seq analysis revealed a progressive increase in fungal biomass during infection (Fig. 3A). Host-derived reads dominated at early stages, accounting for nearly all mapped reads from 0 to 4 dpi, whereas pathogen-derived reads were almost undetectable at 0 dpi and remained below 1% at 3–4 dpi. From 7 dpi onward, the proportion of *Pst*-derived reads increased markedly, reaching approximately 12% at 7 dpi and 40% at 11 dpi, accompanied by a corresponding decline in host-derived reads. These results indicate limited fungal activity during early infection followed by rapid pathogen proliferation at later stages. PCA analysis further revealed clear stage-specific separation of transcriptomes, with pronounced shifts between early infection stages (3–4 dpi) and later stages (7–11 dpi) (Fig. S3A and B). Global expression heatmaps also illustrated progressive transcriptome reprogramming in both barberry and *Pst* throughout the infection course (Fig. S3C and D).

**Fig. 3.**
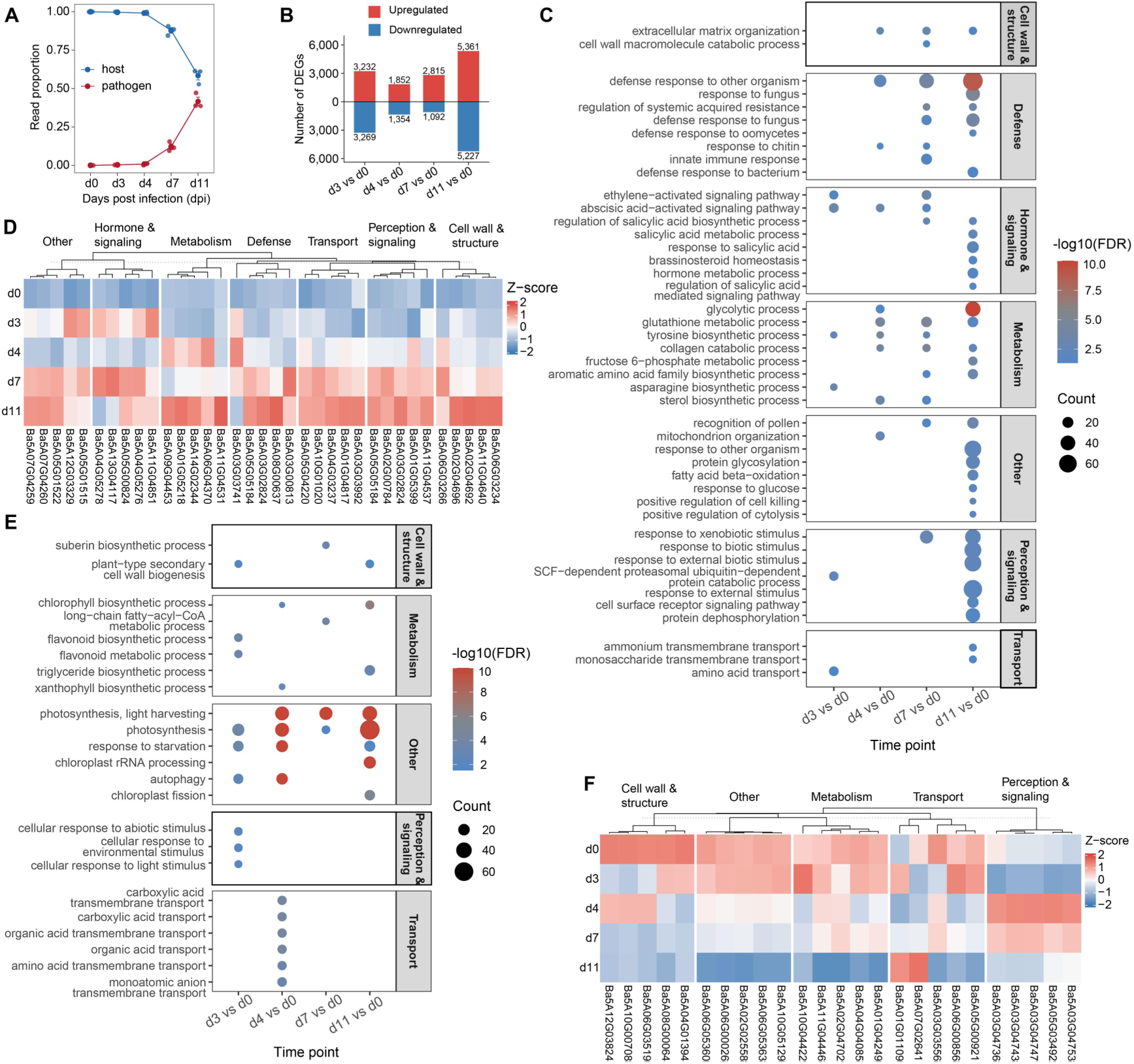
Stage-specific transcriptional reprogramming of barberry during *Puccinia striiformis* f. sp. *tritici* (*Pst*) infection. A, Proportions of reads assigned to the barberry and *Pst* genomes, respectively, across infection stages. Points represent biological replicates; larger points and error bars indicate mean ± SE. B, Numbers of upregulated and downregulated differentially expressed genes (DEGs) at d3, d4, d7 and d11 relative to d0. C, Enriched GO biological process terms among upregulated genes, grouped into broad functional modules, including cell wall and structure, defence, hormone and signalling, metabolism, perception and signalling, transport, and other categories. D, Heatmap of representative upregulated genes selected from the enriched functional modules. E, Enriched GO biological process terms among downregulated genes, grouped into broad functional modules, including metabolism, transport, perception and signalling, cell wall and structure, and other categories. F, Heatmap of representative downregulated genes selected from the enriched functional modules. In the GO bubble plots, bubble size indicates gene count and colour represents enrichment significance as −log10(FDR). In the heatmaps, expression values are shown as row-scaled Z-scores across infection stages.

We next examined host transcriptional responses by comparing infected samples at 3, 4, 7, and 11 dpi with the uninfected control at 0 dpi. The number of differentially expressed genes (DEGs) varied across infection stages and increased sharply at 11 dpi, indicating extensive host transcriptional reprogramming during late infection (Fig. S4A–D and Fig. 3B). Among upregulated genes, GO enrichment analysis revealed a stage-dependent transition from early pathogen perception and signal transduction to later defense activation and metabolic adjustment (Fig. 3C and Table S11). At 3–4 dpi, enriched biological processes were mainly associated with amino acid transport, and asparagine biosynthetic process. At later stages, particularly 7 and 11 dpi, upregulated genes were strongly enriched in defense- and hormone-related processes, including defense response to fungus, regulation of systemic acquired resistance, response to salicylic acid, and salicylic acid-mediated signaling. Processes related to cell wall macromolecule catabolism, and monosaccharide transmembrane transport were also enriched, suggesting substantial structural and metabolic remodeling of host tissues during active infection. KEGG enrichment analysis showed a broadly consistent pattern, with progressive enrichment of signaling- and defense-associated pathways during later infection stages (Fig. S5A). A heatmap of representative upregulated genes further supported these enrichment patterns, showing low basal expression at 0 dpi followed by progressive induction during infection, with many defense-, signaling-, and hormone-associated genes reaching their highest expression at 7–11 dpi, particularly at 11 dpi (Fig. 3D).

In contrast, downregulated host genes were predominantly associated with photosynthesis, plastid function, and primary metabolism. GO enrichment analysis showed strong overrepresentation of terms related to photosynthesis, photosynthetic light harvesting, chloroplast organization, plastid organization, chlorophyll metabolism, and chlorophyll biosynthesis. Several transport-related terms, including amino acid transmembrane transport, organic acid transport, and carboxylic acid transmembrane transport, were also enriched among downregulated genes, particularly at 4 dpi (Fig. 3E and Table S12). In addition, lipid- and secondary metabolism-related processes, including triglyceride biosynthesis, very-long-chain fatty acid biosynthesis, and chlorophyll biosynthesis, were repressed at intermediate and late infection stages. Consistently, KEGG enrichment analysis showed enrichment of photosynthesis- and carbon fixation-related pathways among downregulated genes, especially at intermediate and late stages (Fig. S5B). A heatmap of representative downregulated genes further confirmed this trend, with many photosynthesis-, metabolism-, transport-, and cell wall-related genes showing relatively high expression at 0–3 dpi but progressive repression toward 11 dpi (Fig. 3F).

Pathogen transcriptional activity also changed markedly over the infection course. *Pst* transcriptional reprogramming was limited during the early transition from 3 to 4 dpi, increased substantially from 4 to 7 dpi, and was strongest from 7 to 11 dpi, consistent with the late increase in fungal biomass (Fig. S6A). This transition was accompanied by increased expression of secreted proteins, candidate effectors, CAZymes, proteases, and transporters, with the strongest induction observed at 11 dpi (Fig. S6B). Functional enrichment analysis showed that genes upregulated from 4 to 7 dpi were mainly associated with carbohydrate-, glucan-, and polysaccharide-related metabolic processes, suggesting active nutrient utilization during intermediate colonization (Fig. S6C). Genes induced from 7 to 11 dpi remained enriched in carbohydrate-related processes but additionally showed enrichment in homeostasis-related and developmental regulatory functions, including chemical homeostasis, cellular homeostasis, and regulation of reproductive process. In contrast, genes downregulated from 11 to 7 dpi were enriched in translation-, ribosome biogenesis-, and RNA processing-related processes, suggesting repression of general biosynthetic functions during late colonization (Fig. S6D).

### 3.5. Allele-specific expression dynamic of barberry during *Pst* infection

ASE is widespread in heterozygous diploids and can contribute to diverse biological processes. To investigate its potential involvement in the response of barberry to *Pst* infection, we analyzed haplotype-biased expression in 36,641 one-to-one Ba5A/B alleles across different stages of *Pst* infection. The proportion of ASE genes remained relatively stable across all stages, ranging from approximately 25% to 28%, whereas approximately 27%–29% were classified as unbiased (Fig. 4A). Among the ASE genes, Ba5A- and Ba5B-biased genes were broadly distributed across chromosomes, with no significant difference in their overall numbers (Fig. 4B).

**Fig. 4.**
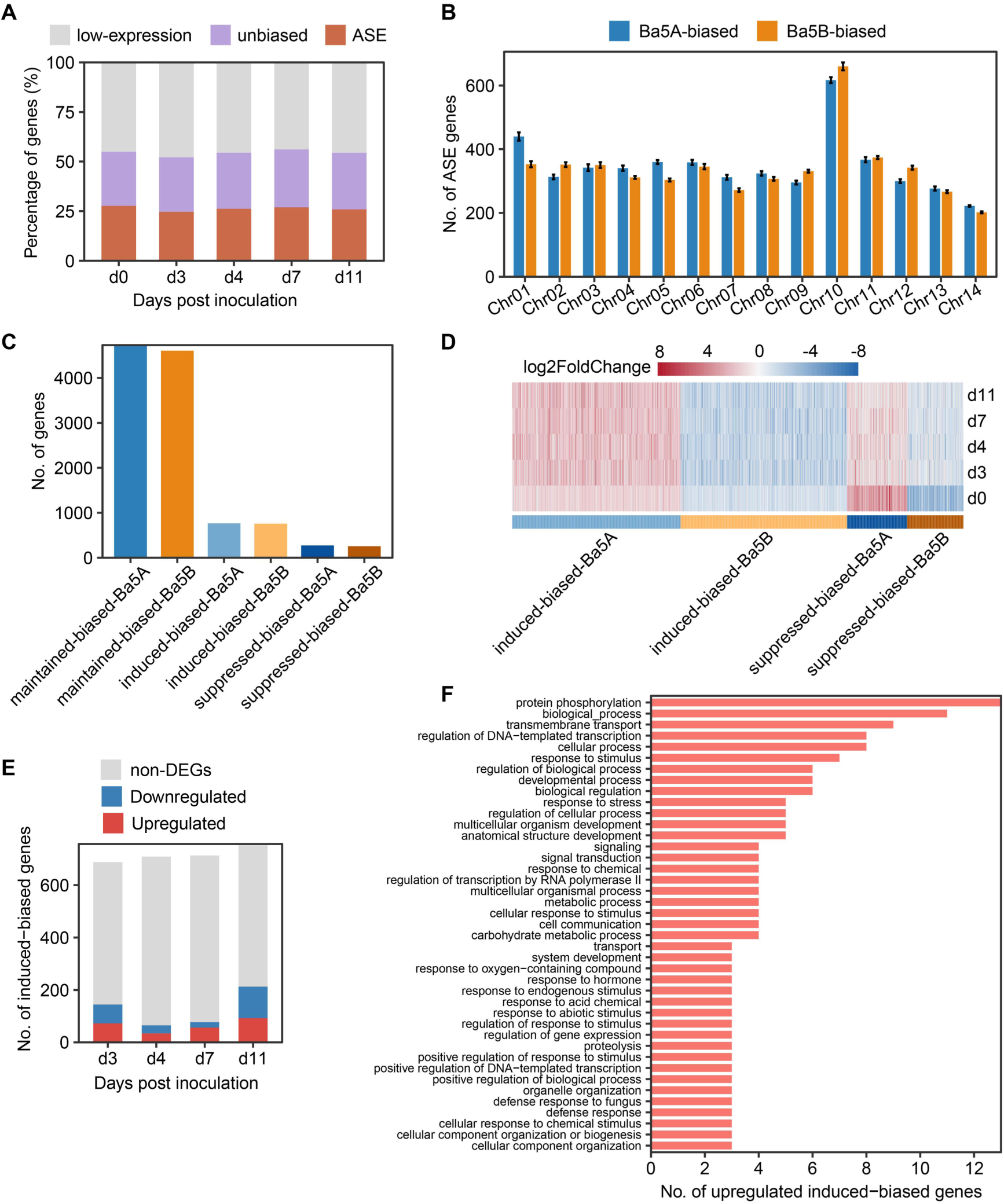
Dynamics of allele-specific expression (ASE) and its association with transcriptional responses during *Puccinia striiformis* f. sp. *tritici* (*Pst*) infection in barberry. A, Proportions of allele expression categories across *Pst* infection stages (0, 3, 4, 7, and 11 days post inoculation). Genes were classified into low-expression, unbiased, and ASE categories. B, Chromosomal distribution of ASE genes across 14 barberry chromosomes, comparing Ba5A- and Ba5B-biased genes. Error bars indicate variability in gene counts across biological replicates. C, Classification of dynamic ASE genes, including maintained, induced, and suppressed ASE genes for both Ba5A and Ba5B haplotypes, highlighting the prevalence of stable and infection-responsive allelic regulation. D, Heatmap of log2FoldChange values for dynamic ASE genes across *Pst* infection stages, illustrating stage-specific transcriptional shifts in induced- and suppressed-biased genes. E, Distribution of induced ASE genes overlapping with differentially expressed genes (DEGs) across *Pst* infection stages, showing proportions of upregulated, downregulated, and non-DEG genes. F, Gene Ontology (GO) biological process annotation of upregulated induced-biased genes.

We next investigated the temporal dynamics of ASE genes during *Pst* infection. Genes that maintained the same haplotype bias from 0 to 11 dpi were labeled as maintained-biased-Ba5A/B, representing the majority of biased genes (Ba5A: 4,729; Ba5B: 4,606), indicating stable haplotype-biased expression across infection stages. Genes that were unbiased at 0 dpi but became biased at subsequent infection stages were labeled as induced-biased-Ba5A/B (Ba5A: 765; Ba5B: 757), reflecting genes whose haplotype-biased expression was infection-induced. And genes that were biased at 0 dpi but lost their bias during infection were labeled as suppressed-biased-Ba5A/B (Ba5A: 273; Ba5B: 256), representing a minor fraction of the genes that were suppressed in their haplotype-biased expression during *Pst* infection (Fig. 4C). Heatmap analysis of log2FoldChange further showed that induced-biased and suppressed-biased genes displayed stage-dependent changes in the magnitude of allelic imbalance (Fig. 4D). These results indicate that ASE genes widely participate in the response to *Pst* infection.

To assess whether infection-responsive allelic bias was accompanied by changes in total gene expression, induced-biased genes were intersected with DEGs identified at each infection stage relative to 0 dpi. We found that a subset of induced-biased genes were also DEGs, including 145 genes at 3 dpi, 65 at 4 dpi, 77 at 7 dpi, and 213 at 11 dpi, suggesting that ASE genes likely play an important role in the response to *Pst* infection (Fig. 4E). Among them, upregulated genes predominated at all stages, especially at 11 dpi, where 121 induced-biased genes were upregulated and 92 were downregulated. These induced-biased DEGs likely represent haplotype-specific contributors to infection-responsive transcriptional reprogramming, with the strongest signal occurring during late infection. We investigated the functional annotation of a non-redundant set of 188 induced-biased genes that were upregulated in at least one infection stage. The functional annotations of these 188 upregulated induced-biased genes were mainly associated with transcriptional regulation, protein phosphorylation and signaling, transmembrane transport, metabolism, and stress- or defense-related responses (Fig. 4F). Notably, several genes were annotated to defense response and defense response to fungus, suggesting that infection-induced haplotype-biased upregulation may contribute to diverse components of host transcriptional reprogramming during *Pst* infection.

## 4. Discussion

Barberry is important not only as a medicinal and ecologically significant shrub, but also as the alternate host for the sexual phase of several cereal rust fungi, including *Pst* (Zhao *et al*. 2016; Sobhani *et al*. 2021). Understanding its genome architecture and interaction with *Pst* is therefore essential for elucidating both barberry biology and wheat stripe rust epidemiology and evolutionary dynamics of *Pst*. Our results show that haplotype-resolved genomic divergence and allele-specific regulatory variation jointly shape the transcriptional landscape of barberry during *Pst* infection. We propose that *Pst* infection elicits a multilayered regulatory response involving both gene-level transcriptional reprogramming and dynamic shifts in ASE, thereby altering the relative contributions of the two haplotypes during host immune responses. This study advances current understanding of plant–pathogen interactions by identifying allele-specific regulation as an underexplored layer of transcriptional control. In long-lived, highly heterozygous woody plants such as barberry, haplotype divergence may provide a flexible regulatory framework that enables fine-tuned responses to biotic and environmental stresses.

Compared with previously available barberry genomes (Bartaula *et al*. 2019; Ruhsam 2024), this study provides, to our knowledge, the first haplotype-resolved chromosome-level genome assembly for barberry, substantially improving the genomic framework for resolving structural heterozygosity, haplotype-specific gene content, and functional divergence between homologous chromosomes. This high-quality reference genome will support a broad range of future studies, including population genomics, germplasm characterization, identification of genes associated with disease resistance and valuable medicinal traits, and marker-assisted improvement of elite barberry varieties. It also provides an important foundation for dissecting the genetic basis of barberry–pathogen interactions and for understanding how haplotype variation contributes to adaptation and phenotypic diversity in this highly heterozygous woody plant.

A prominent feature of the *B. aggregata* genome is the extensive divergence between its two haplotypes. Although most genomic regions remained collinear, a substantial proportion of each haplotype was affected by structural variation or could not be aligned to the corresponding homologous region. These divergent regions were strongly enriched in transposable elements and depleted in genes, indicating that TE accumulation is closely associated with local haplotype differentiation (Vicient and Casacuberta 2017; Emmerson and Catoni 2025). Similar patterns have been reported in other highly heterozygous perennial plants such as *Populus tomentosa* (Li *et al*. 2023), *Vitis retordii* (Liu *et al*. 2024) and *Mimulus guttatus* (Lovell *et al*. 2025), in which repeat-rich and gene-poor regions contribute disproportionately to genome plasticity and structural evolution. In *B. aggregata*, these regions may therefore represent important genomic compartments that promote structural divergence and shape differences in haplotype-specific gene content.

The extensive hemizygosity detected at both the sequence and gene-content levels further supports this interpretation. Rather than representing random gene loss or neutral asymmetry between homologous chromosomes, hemizygosity in *B. aggregata* was associated with non-random functional enrichment, suggesting that the two haplotypes may contribute unequally to genome function. This pattern is consistent with haplotype-resolved studies in other plants showing that homologous chromosomes can differ not only in structure, but also in gene content and functional potential (Cheng *et al*. 2021; Sun *et al*. 2025). For example, in lychee, haplotype-specific structural variation near *CONSTANS*-like genes was associated with fruit maturation differences (Hu *et al*. 2022), and in grape, haplotype-resolved analysis identified a nonsyntenic insertion containing a tandem cluster of *RPP13*-like disease-resistance candidates (Zou *et al*. 2023). More broadly, TE-associated hemizygosity may contribute to intragenomic diversification in barberry and potentially affect development, environmental adaptation, and responses to biotic stress (Peng *et al*. 2025; Pozzi *et al*. 2025).

Dual RNA-seq analysis revealed progressive and coordinated transcriptional reprogramming of both barberry and *Pst* during infection. In the barberry, early infection stages were characterized primarily by activation of pathogen perception and signal transduction pathways, whereas later stages involved a broader physiological transition marked by enhanced defense responses and pronounced repression of photosynthesis and other growth-associated processes, consistent with the growth–defense trade-off and with the central role of chloroplasts in coordinating photosynthetic activity, redox homeostasis, and immune signaling (Huot *et al*. 2014; Lu and Yao 2018). These results suggest that, as fungal colonization progresses, barberry gradually shifts from an early perception- and signaling-dominated response toward a defense-prioritized physiological state (Naidoo *et al*. 2018; Jian *et al*. 2024). *Pst*, by contrast, exhibited a delayed and stage-dependent transcriptional program. The relatively limited transcriptional changes observed at early stages may reflect an initial establishment phase, whereas the subsequent induction of genes encoding secreted proteins, candidate effectors, CAZymes, proteases, and transporters is consistent with increasingly active host manipulation, tissue colonization, and nutrient acquisition (Duplessis *et al*. 2011; Lorrain *et al*. 2019). At the latest infection stage, sustained expression of colonization-associated genes, together with reduced expression of genes involved in translation, ribosome biogenesis, and RNA processing, suggests a transition toward a more specialized late biotrophic program rather than a uniform increase in overall transcriptional activity. Similar stage-specific shifts in pathogenicity-related and core cellular processes have been reported in other rust fungi during host colonization (Lorrain *et al*. 2019; Gay *et al*. 2021; Zhao *et al*. 2021b; Mapuranga *et al*. 2022).

More and more studies have focused on ASE, including its classification, underlying causes, and regulatory mechanisms involved in the formation of complex traits (Cleary and Seoighe 2021; Shi *et al*. 2024). ASE has been reported to influence a range of phenotypic traits, such as fruit color variation (Chen *et al*. 2026) and pathogen resistance (Todesco *et al*. 2010; Wu *et al*. 2026). In this study, we identified a substantial proportion of ASE genes (∼25–28%) across different stages of *Pst* infection in barberry. A subset of these exhibited dynamic regulation, including infection-induced transitions from unbiased expression at 0 dpi to ASE during infection, as well as infection-suppressed transitions from ASE at 0 dpi to unbiased expression during infection. Similar patterns of inconsistent or context-dependent ASE have also been reported in fig fruit peel and pulp, where allelic expression contributions shift between pseudo-haplotypes across tissues (Usai *et al*. 2025). These findings suggest that genomes may dynamically modulate allelic expression in response to developmental or environmental cues, potentially contributing to phenotypic plasticity and heterosis in heterozygous systems.

Notably, a total of 188 induced_bias genes were also identified as DEGs across infection stages relative to 0 dpi, particularly at later infection stages, suggesting coordinated transcriptional regulation at both total gene expression and allele-specific levels. Importantly, a large proportion of induced_bias genes were upregulated during *Pst* infection, indicating preferential activation of specific haplotypes under pathogen pressure. These findings suggest that infection not only modulates overall transcriptional output but also reshapes allelic expression balance, leading to infection-responsive allelic dominance (Moyerbrailean *et al*. 2016). Such patterns highlight allele-specific regulation as an additional layer of transcriptional control in plant immune responses (Wu *et al*. 2026). Functional annotation of these upregulated induced_bias genes also revealed defense-related processes, including fungal defense, suggesting that infection-induced haplotype bias preferentially affects genes involved in immune signaling and stress adaptation.

## 5. Conclusion

This study provides the first haplotype-resolved chromosome-level genome of *B. aggregata* and reveals extensive structural and functional divergence between its two haplotypes. Dual RNA-seq showed coordinated stage-specific transcriptional reprogramming of barberry and *Pst* during infection, while ASE analysis demonstrated that pathogen challenge dynamically reshapes allelic expression balance. These findings establish a genomic framework for studying barberry–rust pathogen interactions and highlight ASE as an additional layer of host immune regulation.

## Acknowledgments

We acknowledge the High-Performance Computing (HPC) of Northwest A&F University for providing computing resources. This work was supported by grants from the National Key R&D Program of China (2024YFD1401000) and China Postdoctoral Science Foundation (2025M783819).

## Declaration of competing interest

The authors declare that they have no conflict of interest.

## Data availability

All raw sequencing data is available at the NCBI database under the BioProjects PRJNA1134302. The assembled genome and gene annotation of barberry are available on Zenodo (https://doi.org/10.5281/zenodo.21550197).

## Supplementary Figure Legends

**Fig. S1.**
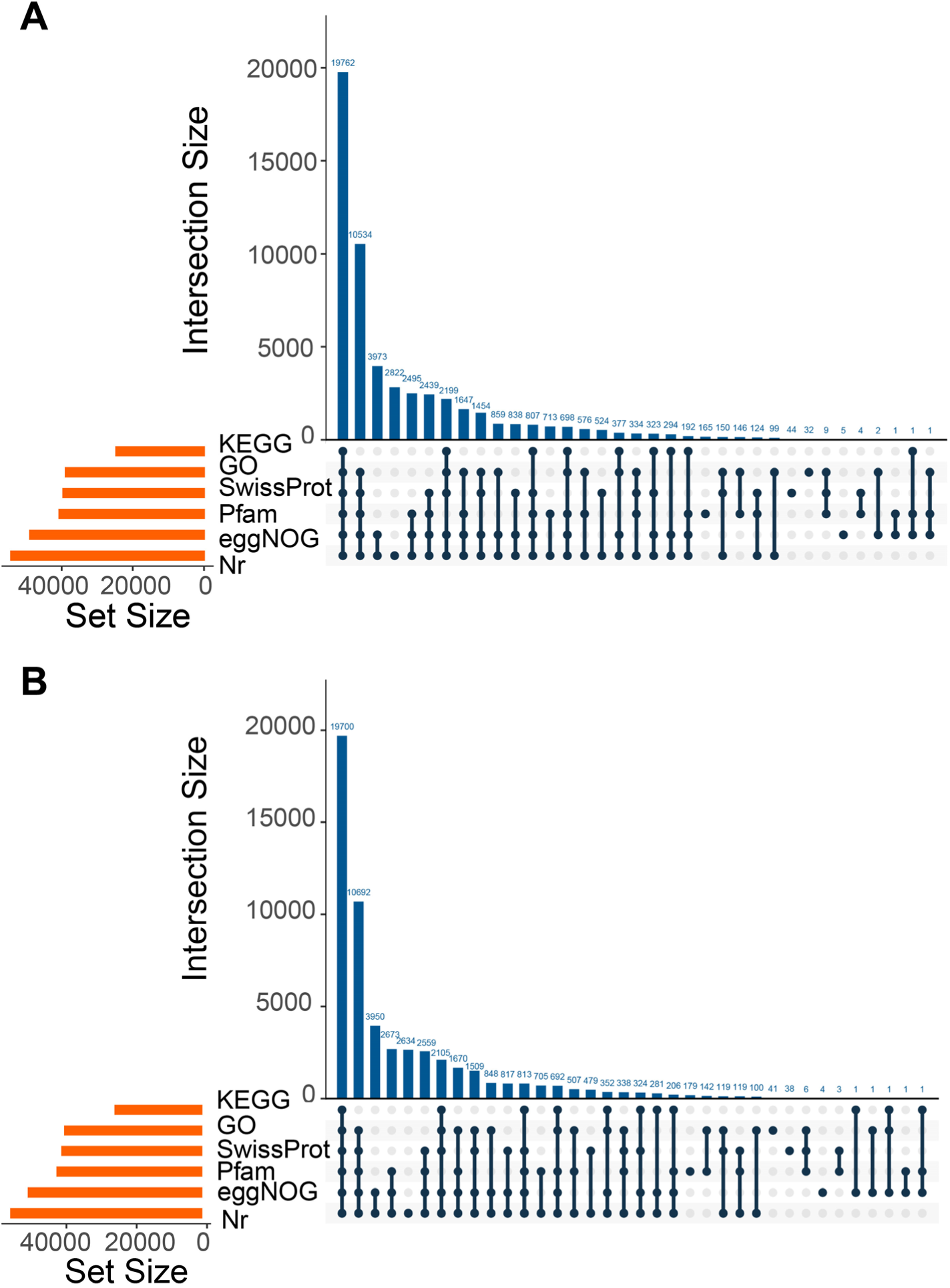
Functional annotation overview of predicted protein-coding genes in the two haplotypes of *Berberis aggregata*. UpSet plots summarize the overlap of functional annotations assigned to predicted proteins in Ba5A (A) and Ba5B (B) across six public databases, including Nr, eggNOG, Pfam, SwissProt, GO and KEGG. The horizontal orange bars on the left indicate the total number of genes annotated in each database, whereas the vertical blue bars represent the sizes of intersections among different annotation combinations. Filled dots connected by lines indicate the corresponding database combinations for each intersection.

**Fig. S2.**
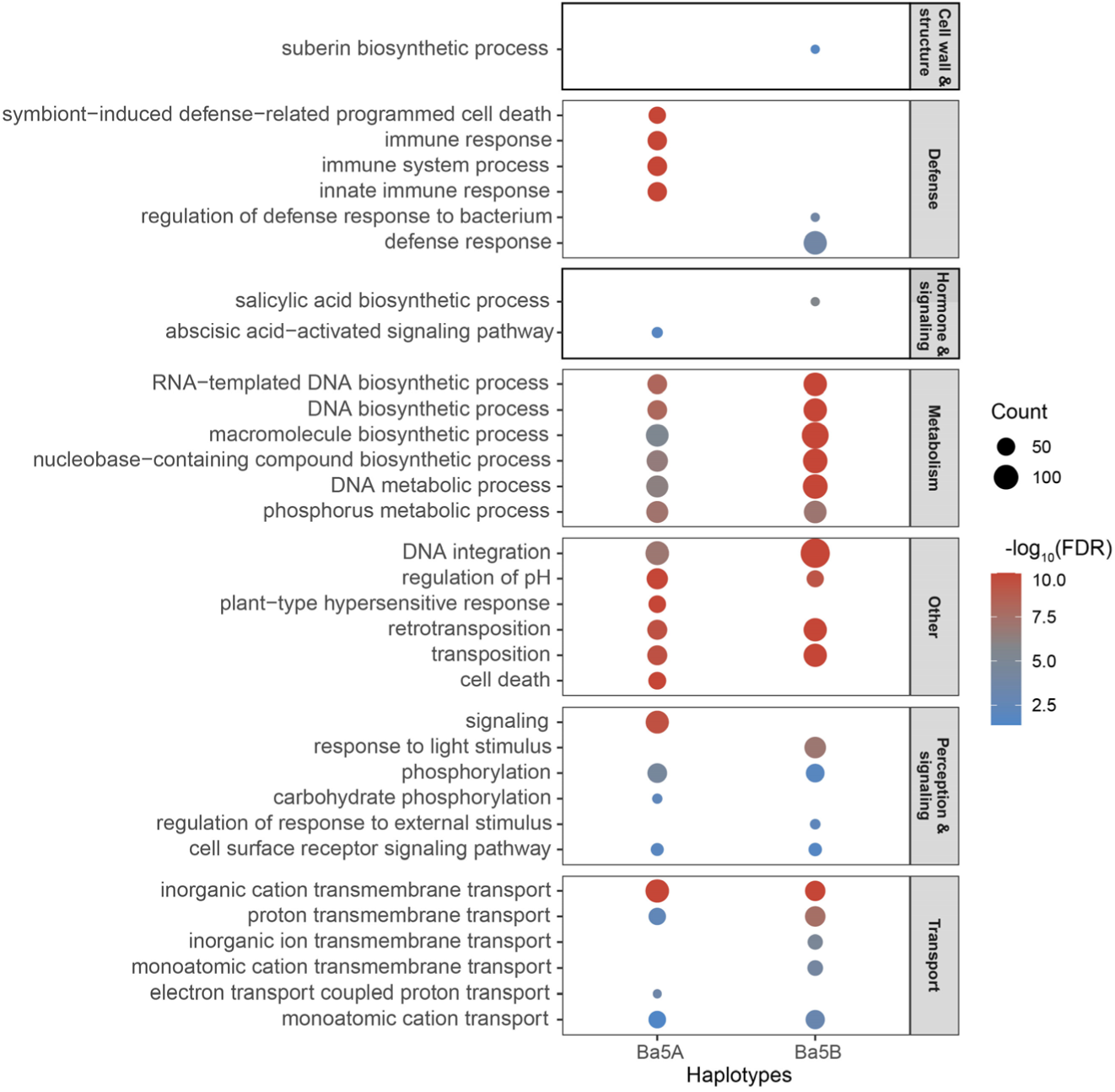
Functional categorization of enriched GO BP terms in haplotype-specific hemizygous genes of Ba5A and Ba5B. Enriched GO biological process terms identified from haplotype-specific hemizygous genes were grouped into manually defined functional modules according to their semantic descriptions. For each module, the six most significant terms ranked by the minimum adjusted *P*-value across Ba5A and Ba5B were retained for plotting. Bubble size denotes gene count and color indicates enrichment significance (−log10(FDR)).

**Fig. S3.**
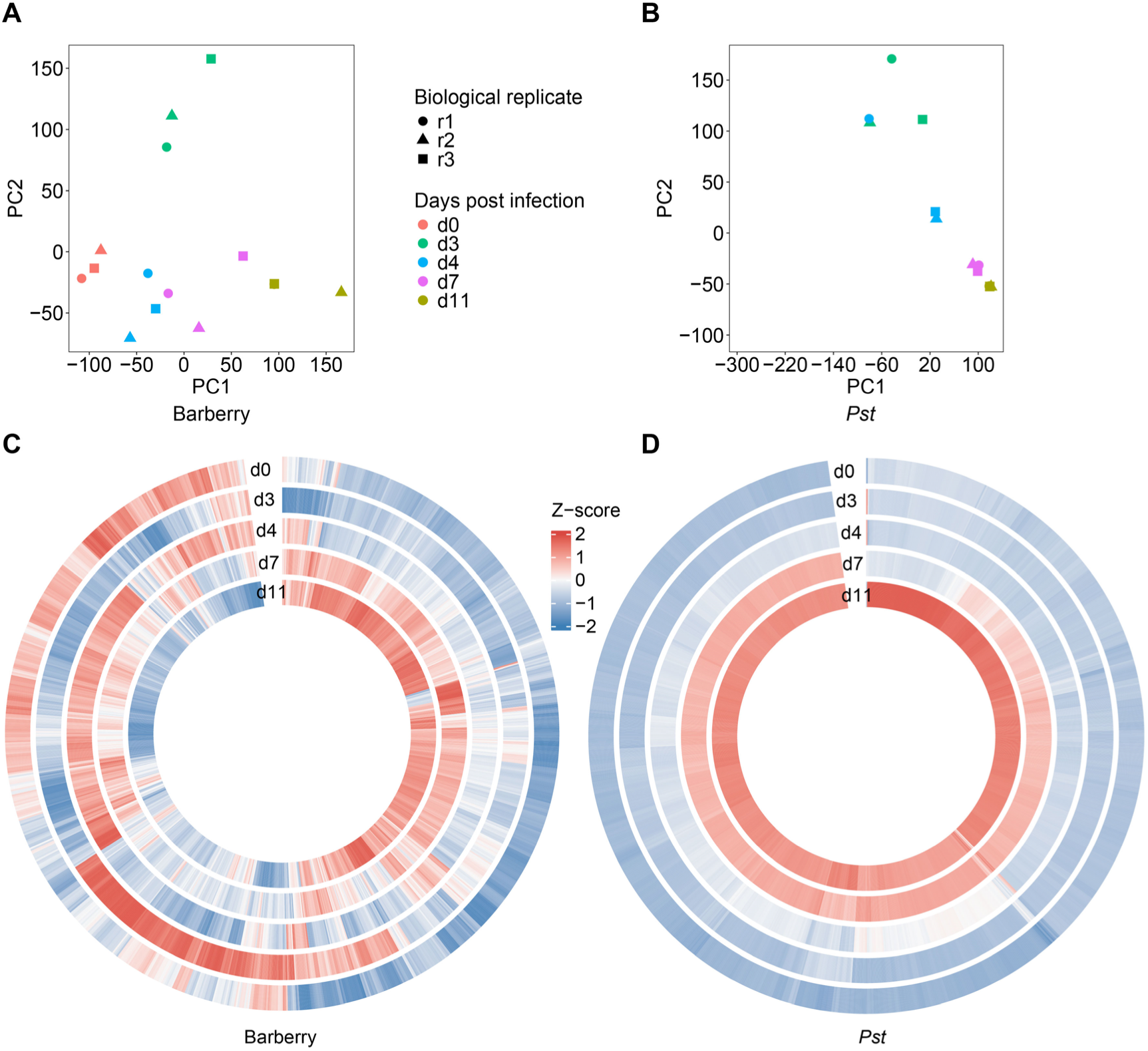
Progressive colonization of barberry by *Puccinia striiformis* f. sp. *Tritici* (*Pst*) revealed by dual RNA-seq. A and B, Principal component analysis reveals stage-specific clustering of barberry and *Pst* transcriptomes. Colors indicate days post infection and shapes indicate biological replicates. The clear separation of samples by stage indicates substantial temporal shifts in global gene expression in both host and pathogen. C and D, Circular heatmaps showing global transcriptome dynamics of barberry and *Pst*, respectively. For each panel, the 1,000 most variably expressed genes were selected, log2-transformed, standardized as row Z-scores, and ordered by hierarchical clustering. Concentric rings represent successive infection stages from outer to inner.

**Fig. S4.**
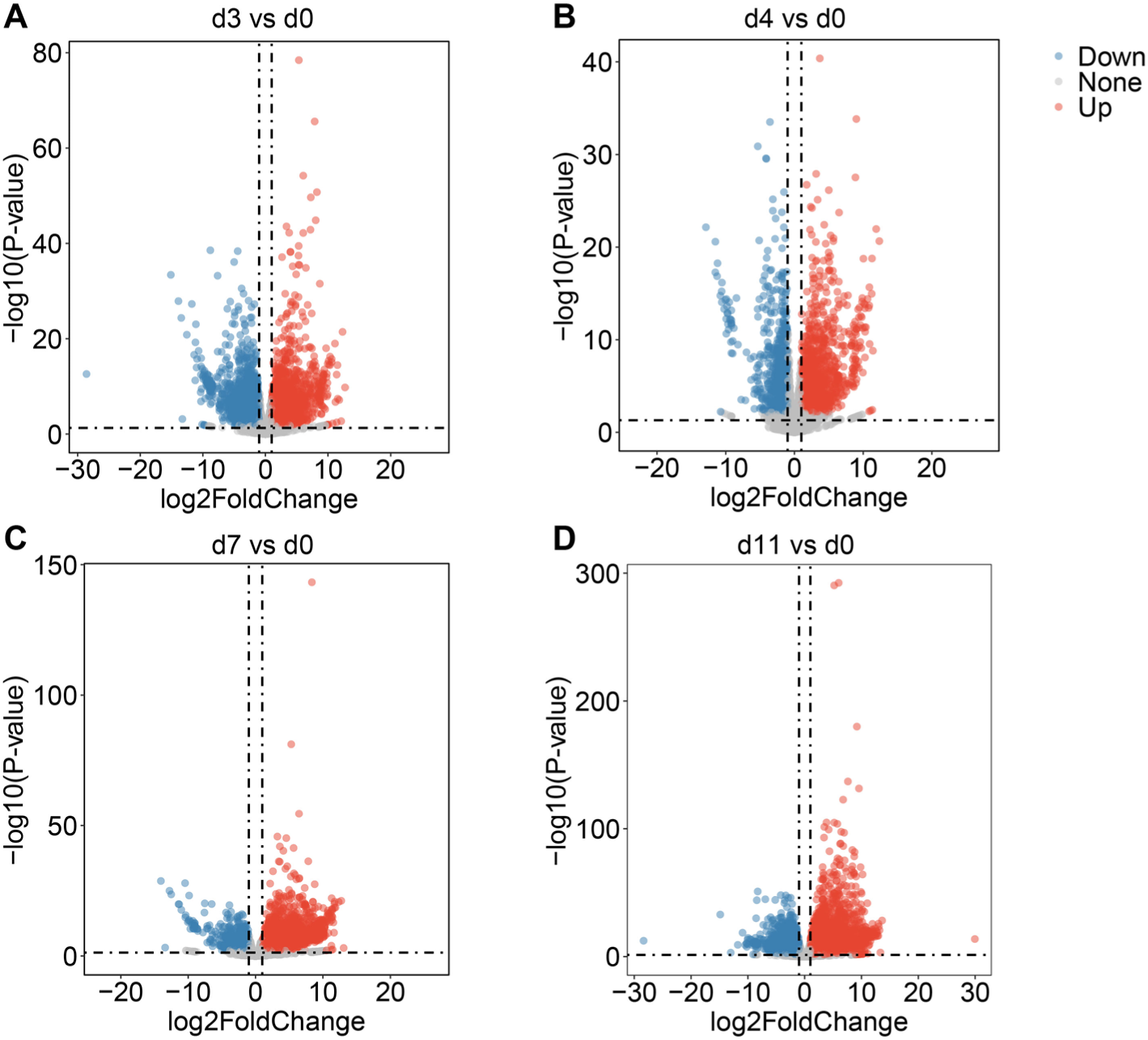
Volcano plots of differentially expressed genes in barberry across infection stages. Volcano plots showing differential gene expression in barberry at d3 vs d0 (A), d4 vs d0 (B), d7 vs d0 (C) and d11 vs d0 (D). Each point represents one gene. The x-axis indicates log2FoldChange, and the y-axis indicates −log10(P-value). Red, blue and grey points denote upregulated, downregulated and non-significant genes, respectively, based on the thresholds used for differential expression analysis.

**Fig. S5.**
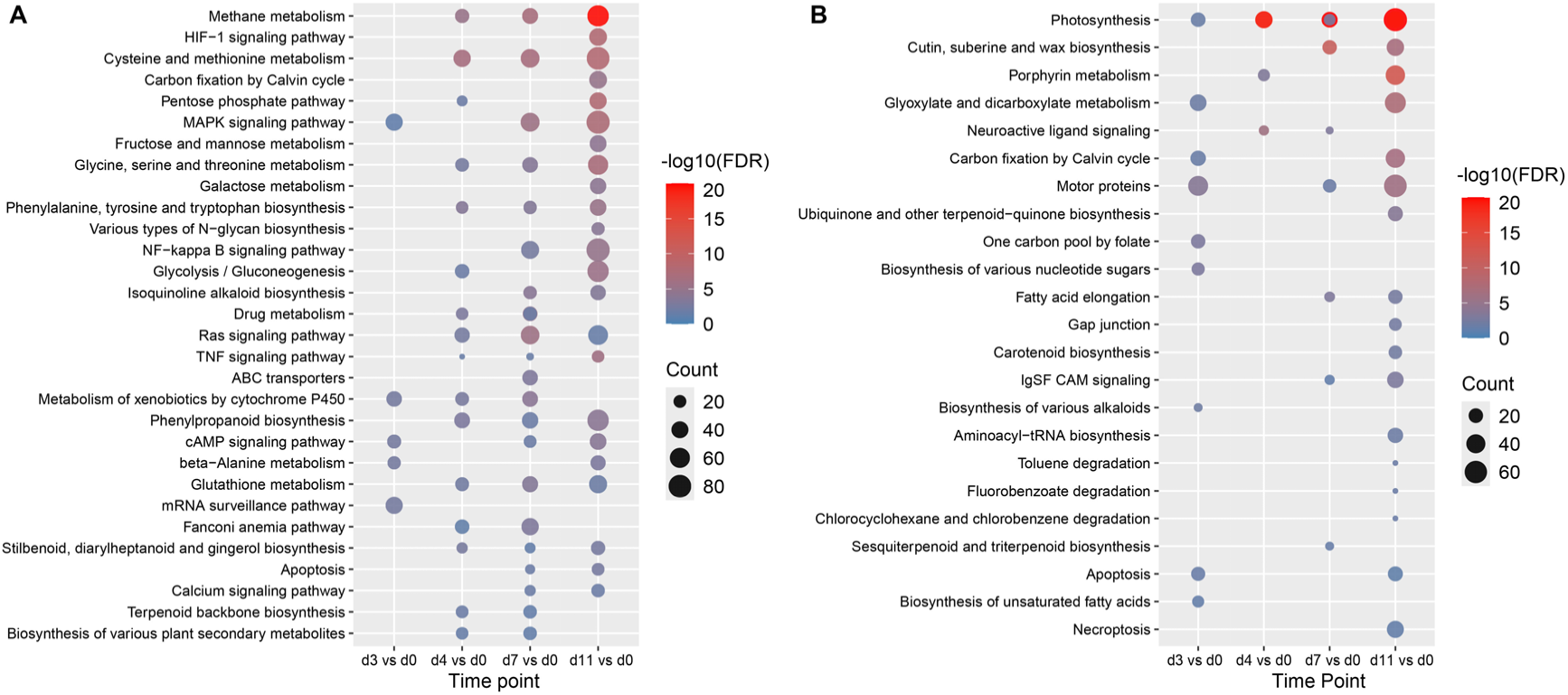
KEGG pathway enrichment analysis of barberry DEGs during *Puccinia striiformis* f. sp. *tritici* (*Pst*) infection. A, KEGG enrichment of upregulated genes at d3 vs d0, d4 vs d0, d7 vs d0 and d11 vs d0. B, KEGG enrichment of downregulated genes at d3 vs d0, d4 vs d0, d7 vs d0 and d11 vs d0. Bubble size indicates the number of genes assigned to each pathway, and bubble colour represents enrichment significance as −log10(FDR). Only significantly enriched pathways are shown.

**Fig. S6.**
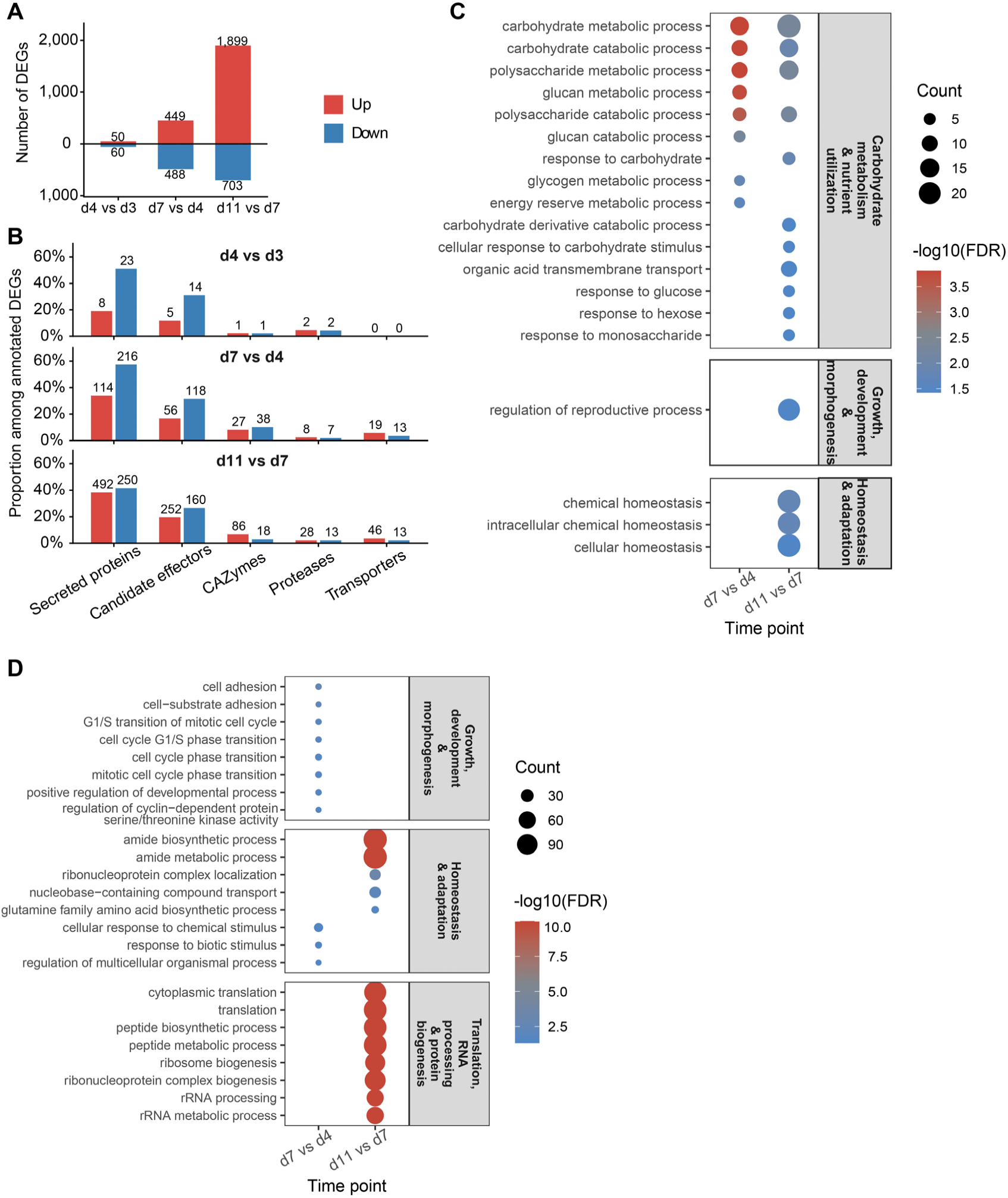
Stage-dependent transcriptional reprogramming of *Puccinia striiformis* f. sp. *tritici* (*Pst*) during colonization of barberry. A, Numbers of upregulated and downregulated differentially expressed genes (DEGs) in *Pst* across successive infection-stage transitions (d4 vs d3, d7 vs d4 and d11 vs d7). B, Numbers of differentially expressed genes belonging to selected virulence- and colonization-associated categories, including secreted proteins, candidate effectors, CAZymes, proteases and transporters, across the three pairwise comparisons. C, Enriched GO biological process terms among upregulated genes, grouped into broad functional modules, including carbohydrate metabolism and nutrient utilization, growth, development and morphogenesis, and homeostasis and adaptation. D, Enriched GO biological process terms among downregulated genes, grouped into broad functional modules, including growth, development and morphogenesis, homeostasis and adaptation, and translation, RNA processing and biogenesis.

## Supplementary Table

Table S1 Summary of sequencing data of *Berberis aggregata* for haplotype-resolved assembly and genome annotation.

Table S2 Statistics of map rate and coverage of different types of sequencing reads.

Table S3 Statistics of Merqury analysis for genome quality assessment.

Table S4 Summary of BUSCO analysis of *Berberis aggregata* genome assembly.

Table S5 Classification of repetitive elements in the *Berberis aggregata* genome assembly.

Table S6 Summary of BUSCO analysis of protein-coding genes in *Berberis aggregata*.

Table S7 Genome annotation statistics.

Table S8 Statistics of variation between two haplotypes.

Table S9 GO biological process enrichment results for all haplotype-specific hemizygous genes in Ba5A and Ba5B.

Table S10 Summary of RNA-seq read mapping to the reference genome across different infection stages.

Table S11 GO biological process enrichment results for upregulated genes in barberry during *Pst* infection.

Table S12 GO biological process enrichment results for downregulated genes in barberry during *Pst* infection.

## Notes

### Competing Interest Statement

The authors have declared no competing interest.

