## Supplementary material for "Haplotype-resolved genome assembly of barberry reveals structural divergence and allele-specific expression during infection by wheat stripe rust pathogen": Table S1-12: Table S1-12.pdf

Table S1 Summary of sequencing data of *Berberis aggregata* for haplotype-resolved assembly and genome annotation.

| Sequencing | Clean base (Gb) | Clean reads | N50 length (bp) | Depth (×) | Sample | Application |
|---|---|---|---|---|---|---|
| HiFi | 97.35 | 5,385,183 | 18,111 | 114.53 | leaves of Ba5 | Genome assembly |
| Hi-C | 100.26 | 668,976,170 | 2 × 150 | 117.95 | leaves of Ba5 | Chromosome construction |
| Illumina | 63.45 | 448,732,915 | 2 × 150 | 74.65 | leaves of Ba5 | Genome evaluation |
| RNA-seq | 3.64 | 24,564,563 | 2 × 150 | — | leaves of Ba5 | Genome annotation |
| RNA-seq | 30.88 | 208,580,233 | 2 × 150 | — | leaves of Ba5 infected by wheat stripe rust pathogen | Genome annotation |

Table S2 Statistics of map rate and coverage of different types of sequencing reads.

| | HiFi DNA | Illumina DNA |
|---|---|---|
| Reads mapped (%) | 99.96 | 99.38 |
| Properly paired (%) | — | 99.03 |
| ≥1× (%) | 100 | 99.94 |
| ≥5× (%) | 99.97 | 99.74 |
| ≥10× (%) | 99.91 | 99.29 |

Table S3 Statistics of Merqury analysis for genome quality assessment.

| Assembly | QV (quality value) | Error rate | Completeness (%) |
|---|---|---|---|
| Ba5A | 57.28 | 1.87E-06 | 87.24 |
| Ba5B | 57.56 | 1.75E-06 | 87.36 |
| both Ba5A and Ba5B | 57.42 | 1.81E-06 | 97.46 |

Table S4 Summary of BUSCO analysis of *Berberis aggregata* genome assembly.

| Statistic | Ba | Haplotype A | Haplotype B |
|---|---|---|---|
| Complete BUSCOs (%) | 1598 (99.0%) | 1597 (98.9%) | 1598 (99.0%) |
| Complete and single-copy BUSCOs (%) | 9 (0.6%) | 1080 (66.9%) | 1091 (67.6%) |
| Complete and duplicated BUSCOs (%) | 1589 (98.5%) | 517 (32.0%) | 507 (31.4%) |
| Fragmented BUSCOs (%) | 6 (0.4%) | 6 (0.4%) | 6 (0.4%) |
| Missing BUSCOs (%) | 10 (0.5%) | 11 (0.7%) | 10 (0.6%) |
| Total BUSCO groups searched | | 1614 | |

Table S5 Classification of repetitive elements in the *Berberis aggregata* genome assembly.

| Category | Ba5A | Ba5B |
|---|---|---|
| LTR/Copia | 109616 (5.90%) | 107378 (5.84%) |
| LTR/Gypsy | 216403 (16.20%) | 195305 (15.93%) |
| LTR/unknown | 180673 (8.37%) | 163249 (6.93%) |
| TIR/CACTA | 224532 (8.86%) | 101773 (3.00%) |
| TIR/Mutator | 173519 (5.15%) | 155950 (15.78%) |
| TIR/PIF_Harbinger | 25093 (0.60%) | 25008 (0.55%) |
| TIR/Tc1_Mariner | 9500 (0.31%) | 10344 (0.31%) |
| TIR/hAT | 223588 (8.36%) | 150897 (5.28%) |
| LINE_element | 2324 (0.09%) | 1745 (0.05%) |
| unknown | 976 (0.05%) | 615 (0.05%) |
| DNA/helitron | 142529 (4.39%) | 172612 (5.91%) |
| repeat_region | 167897 (4.75%) | 165538 (4.57%) |
| Total | 1476650 (63.03%) | 1250414 (64.21%) |

The contents of the cells were counts (percentage of sequence %), LTR, long terminal repeat; TE, transposable element; TIR, terminal inverted repeat.

Table S6 Summary of BUSCO analysis of protein-coding genes in *Berberis aggregata*.

| Statistic | Ba5 | Ba5A | Ba5B |
|---|---|---|---|
| Complete BUSCOs (%) | 1581 (98.0%) | 1536 (95.2%) | 1534 (95.0%) |
| Complete and single-copy BUSCOs (%) | 143 (8.9%) | 1128 (69.9%) | 1124 (69.6%) |
| Complete and duplicated BUSCOs (%) | 1438 (89.1%) | 408 (25.3%) | 410 (25.4%) |
| Fragmented BUSCOs (%) | 16 (1.0%) | 52 (3.2%) | 36 (2.2%) |
| Missing BUSCOs (%) | 17 (1.0%) | 26 (1.6%) | 44 (2.8%) |
| Total BUSCO groups searched | | 1614 | |

Table S7 Genome annotation statistics.

| Parameter | Ba5 | Ba5A | Ba5B |
|---|---|---|---|
| Number of protein-coding genes | 137,127 | 69,793 | 67,334 |
| Number of genes annotated by eggNOG | 98,101 | 48,901 | 49,200 |
| Number of genes annotated by Pfam | 81,654 | 40,648 | 41,006 |
| Number of genes annotated by GO | 77,702 | 38,879 | 38,823 |
| Number of genes annotated by KEGG | 49,129 | 24,653 | 24,476 |
| Number of genes annotated by Nr | 108,425 | 54,233 | 54,192 |
| Number of genes annotated by SwissProt | 79,166 | 39,545 | 39,621 |
| Total number of functionally annotated genes | 109,253 | 54,643 | 54,610 |

Table S8 Statistics of variation between two haplotypes.

| | Variation type | Count | Length of reference (bp) | Length of query (bp) |
|---|---|---|---|---|
| Structural annotations | Syntenic regions | 8,793 | 901,383,859 | 900,055,787 |
| | Inversions | 111 | 30,462,395 | 29,409,138 |
| | Translocations | 2,758 | 24,914,538 | 24,785,165 |
| | Duplications (reference) | 6,706 | 49,371,750 | - |
| | Duplications (query) | 4,855 | - | 33,387,124 |
| | Not aligned (reference) | 10,015 | 149,114,796 | - |
| | Not aligned (query) | 9,805 | - | 184,157,150 |
| Sequence annotations | SNPs | 4,726,951 | 4,726,951 | 4,726,951 |
| | Insertions (INSs and SVs) | 359,209 | - | 3,859,042 |
| | Deletions (DELs and SVs) | 332,586 | 4,810,481 | - |
| | Copygains | 1,655 | - | 9,550,311 |
| | Copylosses | 1,573 | 9,013,274 | - |
| | Highly diverged | 13,048 | 104,853,518 | 102,790,883 |
| | Tandem repeats | 1,578 | 15,238,230 | 15,307,578 |
| | Total | 5,436,600 | 138,642,454 | 136,234,765 |

Table S9 GO biological process enrichment results for all haplotype-specific hemizygous genes in Ba5A and Ba5B.

| ID | Description | GeneRatio | BgRatio | pvalue | p.adjust | qvalue | Count | Ontology | Haplotypes | Category |
|---|---|---|---|---|---|---|---|---|---|---|
| GO:0000281 | mitotic cytokinesis | 9/4261 | 25/38879 | 8.95E-04 | 1.69E-02 | 1.43E-02 | 9 | BP | Ba5A | Other |
| GO:0000481 | maturation of 5S rRNA | 7/4261 | 18/38879 | 2.01E-03 | 2.95E-02 | 2.50E-02 | 7 | BP | Ba5A | Other |
| GO:0000723 | telomere maintenance | 20/4261 | 75/38879 | 1.21E-04 | 3.52E-03 | 2.97E-03 | 20 | BP | Ba5A | Other |
| GO:0000910 | cytokinesis | 9/4261 | 28/38879 | 2.23E-03 | 3.15E-02 | 2.67E-02 | 9 | BP | Ba5A | Other |
| GO:0002376 | immune system process | 60/4261 | 156/38879 | 3.11E-19 | 7.08E-17 | 5.98E-17 | 60 | BP | Ba5A | Defense |
| GO:0005978 | glycogen biosynthetic process | 14/4261 | 49/38879 | 5.75E-04 | 1.20E-02 | 1.01E-02 | 14 | BP | Ba5A | Metabolism |
| GO:0005996 | monosaccharide metabolic process | 12/4261 | 39/38879 | 6.74E-04 | 1.34E-02 | 1.13E-02 | 12 | BP | Ba5A | Metabolism |
| GO:0006082 | organic acid metabolic process | 38/4261 | 213/38879 | 1.75E-03 | 2.69E-02 | 2.27E-02 | 38 | BP | Ba5A | Metabolism |
| GO:0006085 | acetyl-CoA biosynthetic process | 13/4261 | 31/38879 | 9.83E-06 | 4.00E-04 | 3.38E-04 | 13 | BP | Ba5A | Metabolism |
| GO:0006091 | generation of precursor metabolites and energy | 14/4261 | 45/38879 | 2.17E-04 | 5.41E-03 | 4.58E-03 | 14 | BP | Ba5A | Other |
| GO:0006221 | pyrimidine nucleotide biosynthetic process | 11/4261 | 36/38879 | 1.19E-03 | 2.13E-02 | 1.80E-02 | 11 | BP | Ba5A | Metabolism |
| GO:0006259 | DNA metabolic process | 78/4261 | 370/38879 | 1.06E-08 | 6.31E-07 | 5.33E-07 | 78 | BP | Ba5A | Metabolism |
| GO:0006260 | DNA replication | 27/4261 | 137/38879 | 1.80E-03 | 2.75E-02 | 2.32E-02 | 27 | BP | Ba5A | Other |
| GO:0006278 | RNA-templated DNA biosynthetic process | 60/4261 | 229/38879 | 9.01E-11 | 7.93E-09 | 6.70E-09 | 60 | BP | Ba5A | Metabolism |
| GO:0006753 | nucleoside phosphate metabolic process | 13/4261 | 51/38879 | 2.78E-03 | 3.70E-02 | 3.13E-02 | 13 | BP | Ba5A | Metabolism |
| GO:0006793 | phosphorus metabolic process | 77/4261 | 344/38879 | 7.93E-10 | 6.00E-08 | 5.08E-08 | 77 | BP | Ba5A | Metabolism |
| GO:0006796 | phosphate-containing compound metabolic process | 77/4261 | 358/38879 | 5.25E-09 | 3.25E-07 | 2.75E-07 | 77 | BP | Ba5A | Metabolism |
| GO:0006812 | monoatomic cation transport | 46/4261 | 279/38879 | 3.23E-03 | 4.17E-02 | 3.53E-02 | 46 | BP | Ba5A | Transport |
| GO:0006885 | regulation of pH | 72/4261 | 159/38879 | 5.46E-28 | 3.55E-25 | 3.00E-25 | 72 | BP | Ba5A | Regulation |
| GO:0006955 | immune response | 58/4261 | 142/38879 | 3.92E-20 | 1.31E-17 | 1.11E-17 | 58 | BP | Ba5A | Defense |
| GO:0007018 | microtubule-based movement | 40/4261 | 211/38879 | 3.97E-04 | 8.88E-03 | 7.50E-03 | 40 | BP | Ba5A | Other |
| GO:0007052 | mitotic spindle organization | 12/4261 | 43/38879 | 1.75E-03 | 2.69E-02 | 2.27E-02 | 12 | BP | Ba5A | Other |
| GO:0007154 | cell communication | 95/4261 | 432/38879 | 2.52E-11 | 2.75E-09 | 2.33E-09 | 95 | BP | Ba5A | Other |
| GO:0007166 | cell surface receptor signaling pathway | 22/4261 | 96/38879 | 5.94E-04 | 1.23E-02 | 1.04E-02 | 22 | BP | Ba5A | Other |
| GO:0007167 | enzyme-linked receptor protein signaling pathway | 13/4261 | 47/38879 | 1.25E-03 | 2.13E-02 | 1.80E-02 | 13 | BP | Ba5A | Other |

| | | | | | | | | | | |
|---|---|---|---|---|---|---|---|---|---|---|
| GO:0007178 | cell surface receptor protein serine/threonine kinase signaling pathway | 13/4261 | 47/38879 | 1.25E-03 | 2.13E-02 | 1.80E-02 | 13 | BP | Ba5A | Other |
| GO:0007349 | cellularization | 6/4261 | 13/38879 | 1.49E-03 | 2.44E-02 | 2.06E-02 | 6 | BP | Ba5A | Other |
| GO:0008104 | intracellular protein localization | 32/4261 | 179/38879 | 3.72E-03 | 4.65E-02 | 3.93E-02 | 32 | BP | Ba5A | Other |
| GO:0008219 | cell death | 48/4261 | 93/38879 | 3.19E-22 | 1.24E-19 | 1.05E-19 | 48 | BP | Ba5A | Other |
| GO:0009056 | catabolic process | 49/4261 | 295/38879 | 2.08E-03 | 3.01E-02 | 2.54E-02 | 49 | BP | Ba5A | Other |
| GO:0009059 | macromolecule biosynthetic process | 83/4261 | 421/38879 | 8.51E-08 | 4.64E-06 | 3.92E-06 | 83 | BP | Ba5A | Metabolism |
| GO:0009060 | aerobic respiration | 10/4261 | 23/38879 | 7.32E-05 | 2.19E-03 | 1.85E-03 | 10 | BP | Ba5A | Other |
| GO:0009073 | aromatic amino acid family biosynthetic process | 12/4261 | 46/38879 | 3.25E-03 | 4.18E-02 | 3.53E-02 | 12 | BP | Ba5A | Metabolism |
| GO:0009251 | glucan catabolic process | 16/4261 | 42/38879 | 4.26E-06 | 2.04E-04 | 1.72E-04 | 16 | BP | Ba5A | Other |
| GO:0009311 | oligosaccharide metabolic process | 7/4261 | 17/38879 | 1.36E-03 | 2.27E-02 | 1.92E-02 | 7 | BP | Ba5A | Metabolism |
| GO:0009553 | embryo sac development | 19/4261 | 71/38879 | 1.69E-04 | 4.51E-03 | 3.81E-03 | 19 | BP | Ba5A | Development |
| GO:0009561 | megagametogenesis | 13/4261 | 28/38879 | 2.45E-06 | 1.24E-04 | 1.05E-04 | 13 | BP | Ba5A | Other |
| GO:0009607 | response to biotic stimulus | 49/4261 | 298/38879 | 2.57E-03 | 3.49E-02 | 2.95E-02 | 49 | BP | Ba5A | Defense |
| GO:0009626 | plant-type hypersensitive response | 46/4261 | 76/38879 | 2.43E-25 | 1.10E-22 | 9.32E-23 | 46 | BP | Ba5A | Defense |
| GO:0009738 | abscisic acid-activated signaling pathway | 15/4261 | 56/38879 | 7.85E-04 | 1.53E-02 | 1.29E-02 | 15 | BP | Ba5A | Other |
| GO:0009767 | photosynthetic electron transport chain | 13/4261 | 40/38879 | 2.22E-04 | 5.46E-03 | 4.61E-03 | 13 | BP | Ba5A | Transport |
| GO:0009890 | negative regulation of biosynthetic process | 26/4261 | 128/38879 | 1.38E-03 | 2.29E-02 | 1.93E-02 | 26 | BP | Ba5A | Metabolism |
| GO:0009892 | negative regulation of metabolic process | 39/4261 | 211/38879 | 7.76E-04 | 1.52E-02 | 1.29E-02 | 39 | BP | Ba5A | Metabolism |
| GO:0009893 | positive regulation of metabolic process | 54/4261 | 274/38879 | 1.43E-05 | 5.64E-04 | 4.77E-04 | 54 | BP | Ba5A | Metabolism |
| GO:0009894 | regulation of catabolic process | 17/4261 | 58/38879 | 1.09E-04 | 3.23E-03 | 2.73E-03 | 17 | BP | Ba5A | Regulation |
| GO:0009896 | positive regulation of catabolic process | 10/4261 | 34/38879 | 2.71E-03 | 3.63E-02 | 3.07E-02 | 10 | BP | Ba5A | Regulation |
| GO:0010467 | gene expression | 38/4261 | 215/38879 | 2.08E-03 | 3.01E-02 | 2.54E-02 | 38 | BP | Ba5A | Other |
| GO:0010558 | negative regulation of macromolecule biosynthetic process | 26/4261 | 126/38879 | 1.09E-03 | 1.98E-02 | 1.67E-02 | 26 | BP | Ba5A | Metabolism |
| GO:0010604 | positive regulation of macromolecule metabolic process | 45/4261 | 241/38879 | 2.56E-04 | 6.07E-03 | 5.13E-03 | 45 | BP | Ba5A | Metabolism |

| | | | | | | | | | | |
|---|---|---|---|---|---|---|---|---|---|---|
| GO:0010605 | negative regulation of macromolecule metabolic process | 39/4261 | 195/38879 | 1.49E-04 | 4.16E-03 | 3.52E-03 | 39 | BP | Ba5A | Metabolism |
| GO:0010608 | post-transcriptional regulation of gene expression | 20/4261 | 84/38879 | 6.16E-04 | 1.26E-02 | 1.07E-02 | 20 | BP | Ba5A | Regulation |
| GO:0010638 | positive regulation of organelle organization | 7/4261 | 19/38879 | 2.88E-03 | 3.82E-02 | 3.23E-02 | 7 | BP | Ba5A | Regulation |
| GO:0012501 | programmed cell death | 48/4261 | 102/38879 | 5.01E-20 | 1.37E-17 | 1.16E-17 | 48 | BP | Ba5A | Other |
| GO:0015074 | DNA integration | 90/4261 | 432/38879 | 1.53E-09 | 1.10E-07 | 9.29E-08 | 90 | BP | Ba5A | Other |
| GO:0015833 | peptide transport | 27/4261 | 119/38879 | 1.81E-04 | 4.70E-03 | 3.97E-03 | 27 | BP | Ba5A | Transport |
| GO:0015936 | coenzyme A metabolic process | 13/4261 | 52/38879 | 3.35E-03 | 4.28E-02 | 3.62E-02 | 13 | BP | Ba5A | Metabolism |
| GO:0015990 | electron transport coupled proton transport | 8/4261 | 12/38879 | 6.81E-06 | 2.95E-04 | 2.49E-04 | 8 | BP | Ba5A | Transport |
| GO:0016104 | triterpenoid biosynthetic process | 18/4261 | 39/38879 | 3.20E-08 | 1.86E-06 | 1.57E-06 | 18 | BP | Ba5A | Metabolism |
| GO:0016310 | phosphorylation | 60/4261 | 287/38879 | 6.68E-07 | 3.50E-05 | 2.96E-05 | 60 | BP | Ba5A | Other |
| GO:0019637 | organophosphate metabolic process | 19/4261 | 77/38879 | 5.20E-04 | 1.09E-02 | 9.22E-03 | 19 | BP | Ba5A | Metabolism |
| GO:0019684 | photosynthesis, light reaction | 17/4261 | 47/38879 | 4.92E-06 | 2.27E-04 | 1.92E-04 | 17 | BP | Ba5A | Other |
| GO:0019856 | pyrimidine nucleobase biosynthetic process | 8/4261 | 24/38879 | 3.02E-03 | 3.92E-02 | 3.32E-02 | 8 | BP | Ba5A | Metabolism |
| GO:0022607 | cellular component assembly | 43/4261 | 215/38879 | 7.08E-05 | 2.15E-03 | 1.81E-03 | 43 | BP | Ba5A | Other |
| GO:0023052 | signaling | 87/4261 | 367/38879 | 2.59E-12 | 3.53E-10 | 2.99E-10 | 87 | BP | Ba5A | Other |
| GO:0031399 | regulation of protein modification process | 17/4261 | 72/38879 | 1.69E-03 | 2.63E-02 | 2.22E-02 | 17 | BP | Ba5A | Regulation |
| GO:0031401 | positive regulation of protein modification process | 13/4261 | 49/38879 | 1.89E-03 | 2.86E-02 | 2.42E-02 | 13 | BP | Ba5A | Regulation |
| GO:0032196 | transposition | 62/4261 | 224/38879 | 3.50E-12 | 4.15E-10 | 3.51E-10 | 62 | BP | Ba5A | Other |
| GO:0032197 | retrotransposition | 62/4261 | 224/38879 | 3.50E-12 | 4.15E-10 | 3.51E-10 | 62 | BP | Ba5A | Other |
| GO:0032787 | monocarboxylic acid metabolic process | 24/4261 | 99/38879 | 1.38E-04 | 3.89E-03 | 3.29E-03 | 24 | BP | Ba5A | Metabolism |
| GO:0033013 | tetrapyrrole metabolic process | 6/4261 | 11/38879 | 4.89E-04 | 1.05E-02 | 8.87E-03 | 6 | BP | Ba5A | Metabolism |
| GO:0033036 | macromolecule localization | 39/4261 | 209/38879 | 6.41E-04 | 1.30E-02 | 1.10E-02 | 39 | BP | Ba5A | Other |
| GO:0033554 | cellular response to stress | 77/4261 | 305/38879 | 1.64E-12 | 2.36E-10 | 1.99E-10 | 77 | BP | Ba5A | Defense |
| GO:0033865 | nucleoside bisphosphate metabolic process | 5/4261 | 11/38879 | 4.12E-03 | 4.93E-02 | 4.17E-02 | 5 | BP | Ba5A | Metabolism |
| GO:0033875 | ribonucleoside bisphosphate metabolic process | 5/4261 | 11/38879 | 4.12E-03 | 4.93E-02 | 4.17E-02 | 5 | BP | Ba5A | Metabolism |

| | | | | | | | | | | |
|---|---|---|---|---|---|---|---|---|---|---|
| GO:0034032 | purine nucleoside bisphosphate metabolic process | 5/4261 | 11/38879 | 4.12E-03 | 4.93E-02 | 4.17E-02 | 5 | BP | Ba5A | Metabolism |
| GO:0034050 | symbiont-induced defense-related programmed cell death | 45/4261 | 66/38879 | 4.23E-28 | 3.55E-25 | 3.00E-25 | 45 | BP | Ba5A | Defense |
| GO:0034248 | regulation of amide metabolic process | 8/4261 | 23/38879 | 2.22E-03 | 3.15E-02 | 2.67E-02 | 8 | BP | Ba5A | Metabolism |
| GO:0034654 | nucleobase-containing compound biosynthetic process | 72/4261 | 325/38879 | 4.40E-09 | 2.85E-07 | 2.41E-07 | 72 | BP | Ba5A | Metabolism |
| GO:0042886 | amide transport | 27/4261 | 121/38879 | 2.42E-04 | 5.83E-03 | 4.93E-03 | 27 | BP | Ba5A | Transport |
| GO:0043207 | response to external biotic stimulus | 49/4261 | 298/38879 | 2.57E-03 | 3.49E-02 | 2.95E-02 | 49 | BP | Ba5A | Defense |
| GO:0043436 | oxoacid metabolic process | 38/4261 | 213/38879 | 1.75E-03 | 2.69E-02 | 2.27E-02 | 38 | BP | Ba5A | Metabolism |
| GO:0043487 | regulation of RNA stability | 6/4261 | 11/38879 | 4.89E-04 | 1.05E-02 | 8.87E-03 | 6 | BP | Ba5A | Regulation |
| GO:0044085 | cellular component biogenesis | 60/4261 | 301/38879 | 3.40E-06 | 1.65E-04 | 1.40E-04 | 60 | BP | Ba5A | Other |
| GO:0044281 | small molecule metabolic process | 61/4261 | 326/38879 | 2.20E-05 | 7.63E-04 | 6.45E-04 | 61 | BP | Ba5A | Metabolism |
| GO:0044550 | secondary metabolite biosynthetic process | 18/4261 | 70/38879 | 4.23E-04 | 9.37E-03 | 7.92E-03 | 18 | BP | Ba5A | Metabolism |
| GO:0045087 | innate immune response | 58/4261 | 164/38879 | 1.17E-16 | 2.28E-14 | 1.92E-14 | 58 | BP | Ba5A | Defense |
| GO:0045184 | establishment of protein localization | 27/4261 | 118/38879 | 1.56E-04 | 4.29E-03 | 3.63E-03 | 27 | BP | Ba5A | Other |
| GO:0045292 | mRNA cis splicing, via spliceosome | 13/4261 | 46/38879 | 1.00E-03 | 1.85E-02 | 1.56E-02 | 13 | BP | Ba5A | Other |
| GO:0045934 | negative regulation of nucleobase-containing compound metabolic process | 26/4261 | 134/38879 | 2.72E-03 | 3.63E-02 | 3.07E-02 | 26 | BP | Ba5A | Metabolism |
| GO:0046294 | formaldehyde catabolic process | 14/4261 | 56/38879 | 2.37E-03 | 3.30E-02 | 2.79E-02 | 14 | BP | Ba5A | Other |
| GO:0046835 | carbohydrate phosphorylation | 12/4261 | 35/38879 | 2.18E-04 | 5.41E-03 | 4.58E-03 | 12 | BP | Ba5A | Other |
| GO:0046907 | intracellular transport | 30/4261 | 146/38879 | 5.10E-04 | 1.09E-02 | 9.19E-03 | 30 | BP | Ba5A | Transport |
| GO:0048518 | positive regulation of biological process | 68/4261 | 427/38879 | 1.07E-03 | 1.96E-02 | 1.66E-02 | 68 | BP | Ba5A | Regulation |
| GO:0048519 | negative regulation of biological process | 55/4261 | 349/38879 | 3.78E-03 | 4.68E-02 | 3.96E-02 | 55 | BP | Ba5A | Regulation |
| GO:0048522 | positive regulation of cellular process | 57/4261 | 338/38879 | 6.63E-04 | 1.33E-02 | 1.12E-02 | 57 | BP | Ba5A | Regulation |
| GO:0048523 | negative regulation of cellular process | 46/4261 | 272/38879 | 1.94E-03 | 2.93E-02 | 2.48E-02 | 46 | BP | Ba5A | Regulation |
| GO:0051130 | positive regulation of cellular component organization | 11/4261 | 35/38879 | 9.20E-04 | 1.73E-02 | 1.46E-02 | 11 | BP | Ba5A | Regulation |
| GO:0051246 | regulation of protein metabolic process | 33/4261 | 126/38879 | 1.42E-06 | 7.30E-05 | 6.17E-05 | 33 | BP | Ba5A | Metabolism |

| | | | | | | | | | | |
|---|---|---|---|---|---|---|---|---|---|---|
| GO:0051247 | positive regulation of protein metabolic process | 22/4261 | 78/38879 | 2.21E-05 | 7.63E-04 | 6.45E-04 | 22 | BP | Ba5A | Metabolism |
| GO:0051248 | negative regulation of protein metabolic process | 13/4261 | 38/38879 | 1.23E-04 | 3.54E-03 | 2.99E-03 | 13 | BP | Ba5A | Metabolism |
| GO:0051253 | negative regulation of RNA metabolic process | 26/4261 | 129/38879 | 1.56E-03 | 2.53E-02 | 2.14E-02 | 26 | BP | Ba5A | Metabolism |
| GO:0051641 | cellular localization | 36/4261 | 179/38879 | 2.34E-04 | 5.70E-03 | 4.82E-03 | 36 | BP | Ba5A | Other |
| GO:0051649 | establishment of localization in cell | 30/4261 | 142/38879 | 3.11E-04 | 7.07E-03 | 5.98E-03 | 30 | BP | Ba5A | Other |
| GO:0051707 | response to other organism | 49/4261 | 297/38879 | 2.40E-03 | 3.30E-02 | 2.79E-02 | 49 | BP | Ba5A | Defense |
| GO:0055086 | nucleobase-containing small molecule metabolic process | 16/4261 | 67/38879 | 2.01E-03 | 2.95E-02 | 2.50E-02 | 16 | BP | Ba5A | Metabolism |
| GO:0061640 | cytoskeleton-dependent cytokinesis | 9/4261 | 28/38879 | 2.23E-03 | 3.15E-02 | 2.67E-02 | 9 | BP | Ba5A | Other |
| GO:0065009 | regulation of molecular function | 24/4261 | 118/38879 | 2.03E-03 | 2.96E-02 | 2.50E-02 | 24 | BP | Ba5A | Regulation |
| GO:0070475 | rRNA base methylation | 14/4261 | 53/38879 | 1.34E-03 | 2.26E-02 | 1.91E-02 | 14 | BP | Ba5A | Other |
| GO:0070925 | organelle assembly | 14/4261 | 48/38879 | 4.56E-04 | 9.95E-03 | 8.41E-03 | 14 | BP | Ba5A | Other |
| GO:0071277 | cellular response to calcium ion | 8/4261 | 24/38879 | 3.02E-03 | 3.92E-02 | 3.32E-02 | 8 | BP | Ba5A | Defense |
| GO:0071897 | DNA biosynthetic process | 60/4261 | 230/38879 | 1.09E-10 | 9.29E-09 | 7.85E-09 | 60 | BP | Ba5A | Metabolism |
| GO:0080134 | regulation of response to stress | 26/4261 | 135/38879 | 3.02E-03 | 3.92E-02 | 3.32E-02 | 26 | BP | Ba5A | Defense |
| GO:0098542 | defense response to other organism | 56/4261 | 313/38879 | 1.61E-04 | 4.40E-03 | 3.72E-03 | 56 | BP | Ba5A | Defense |
| GO:0098662 | inorganic cation transmembrane transport | 90/4261 | 235/38879 | 6.01E-28 | 3.55E-25 | 3.00E-25 | 90 | BP | Ba5A | Transport |
| GO:0140014 | mitotic nuclear division | 8/4261 | 24/38879 | 3.02E-03 | 3.92E-02 | 3.32E-02 | 8 | BP | Ba5A | Other |
| GO:1902600 | proton transmembrane transport | 45/4261 | 234/38879 | 1.28E-04 | 3.64E-03 | 3.07E-03 | 45 | BP | Ba5A | Transport |
| GO:1902850 | microtubule cytoskeleton organization involved in mitosis | 8/4261 | 15/38879 | 6.53E-05 | 2.02E-03 | 1.71E-03 | 8 | BP | Ba5A | Other |
| GO:1903311 | regulation of mRNA metabolic process | 9/4261 | 23/38879 | 4.39E-04 | 9.65E-03 | 8.15E-03 | 9 | BP | Ba5A | Metabolism |
| GO:2001252 | positive regulation of chromosome organization | 7/4261 | 18/38879 | 2.01E-03 | 2.95E-02 | 2.50E-02 | 7 | BP | Ba5A | Regulation |
| GO:0000470 | maturation of LSU-rRNA | 9/4522 | 29/38823 | 4.43E-03 | 4.23E-02 | 3.56E-02 | 9 | BP | Ba5B | Other |
| GO:0001101 | response to acid chemical | 65/4522 | 399/38823 | 3.36E-03 | 3.31E-02 | 2.78E-02 | 65 | BP | Ba5B | Defense |
| GO:0001676 | long-chain fatty acid metabolic process | 7/4522 | 17/38823 | 1.94E-03 | 2.14E-02 | 1.80E-02 | 7 | BP | Ba5B | Metabolism |
| GO:0002182 | cytoplasmic translational elongation | 6/4522 | 12/38823 | 1.23E-03 | 1.59E-02 | 1.34E-02 | 6 | BP | Ba5B | Other |
| GO:0002831 | regulation of response to biotic stimulus | 12/4522 | 38/38823 | 8.99E-04 | 1.27E-02 | 1.06E-02 | 12 | BP | Ba5B | Defense |

| | | | | | | | | | | |
|---|---|---|---|---|---|---|---|---|---|---|
| GO:0006082 | organic acid metabolic process | 43/4522 | 220/38823 | 4.58E-04 | 7.45E-03 | 6.27E-03 | 43 | BP | Ba5B | Metabolism |
| GO:0006090 | pyruvate metabolic process | 17/4522 | 54/38823 | 8.65E-05 | 1.78E-03 | 1.50E-03 | 17 | BP | Ba5B | Metabolism |
| GO:0006091 | generation of precursor metabolites and energy | 17/4522 | 52/38823 | 5.06E-05 | 1.17E-03 | 9.84E-04 | 17 | BP | Ba5B | Other |
| GO:0006259 | DNA metabolic process | 101/4522 | 368/38823 | 8.18E-17 | 6.95E-15 | 5.84E-15 | 101 | BP | Ba5B | Metabolism |
| GO:0006278 | RNA-templated DNA biosynthetic process | 89/4522 | 246/38823 | 1.02E-23 | 3.57E-21 | 3.00E-21 | 89 | BP | Ba5B | Metabolism |
| GO:0006364 | rRNA processing | 36/4522 | 187/38823 | 1.67E-03 | 1.92E-02 | 1.62E-02 | 36 | BP | Ba5B | Other |
| GO:0006778 | porphyrin-containing compound metabolic process | 9/4522 | 19/38823 | 1.21E-04 | 2.36E-03 | 1.99E-03 | 9 | BP | Ba5B | Metabolism |
| GO:0006779 | porphyrin-containing compound biosynthetic process | 8/4522 | 19/38823 | 7.74E-04 | 1.13E-02 | 9.50E-03 | 8 | BP | Ba5B | Metabolism |
| GO:0006787 | porphyrin-containing compound catabolic process | 8/4522 | 17/38823 | 3.09E-04 | 5.27E-03 | 4.43E-03 | 8 | BP | Ba5B | Other |
| GO:0006793 | phosphorus metabolic process | 83/4522 | 365/38823 | 1.48E-09 | 9.67E-08 | 8.13E-08 | 83 | BP | Ba5B | Metabolism |
| GO:0006796 | phosphate-containing compound metabolic process | 85/4522 | 378/38823 | 1.68E-09 | 1.05E-07 | 8.79E-08 | 85 | BP | Ba5B | Metabolism |
| GO:0006811 | monoatomic ion transport | 57/4522 | 308/38823 | 2.77E-04 | 4.91E-03 | 4.13E-03 | 57 | BP | Ba5B | Transport |
| GO:0006812 | monoatomic cation transport | 57/4522 | 287/38823 | 3.86E-05 | 9.25E-04 | 7.78E-04 | 57 | BP | Ba5B | Transport |
| GO:0006873 | intracellular monoatomic ion homeostasis | 16/4522 | 55/38823 | 3.82E-04 | 6.40E-03 | 5.39E-03 | 16 | BP | Ba5B | Other |
| GO:0006885 | regulation of pH | 44/4522 | 127/38823 | 9.61E-12 | 6.90E-10 | 5.81E-10 | 44 | BP | Ba5B | Regulation |
| GO:0006952 | defense response | 87/4522 | 466/38823 | 5.97E-06 | 1.69E-04 | 1.42E-04 | 87 | BP | Ba5B | Defense |
| GO:0006970 | response to osmotic stress | 37/4522 | 194/38823 | 1.72E-03 | 1.98E-02 | 1.66E-02 | 37 | BP | Ba5B | Defense |
| GO:0007154 | cell communication | 67/4522 | 419/38823 | 4.59E-03 | 4.37E-02 | 3.68E-02 | 67 | BP | Ba5B | Other |
| GO:0007166 | cell surface receptor signaling pathway | 24/4522 | 111/38823 | 1.94E-03 | 2.14E-02 | 1.80E-02 | 24 | BP | Ba5B | Other |
| GO:0009056 | catabolic process | 63/4522 | 327/38823 | 4.10E-05 | 9.74E-04 | 8.19E-04 | 63 | BP | Ba5B | Other |
| GO:0009057 | macromolecule catabolic process | 40/4522 | 228/38823 | 5.42E-03 | 4.96E-02 | 4.17E-02 | 40 | BP | Ba5B | Other |
| GO:0009059 | macromolecule biosynthetic process | 121/4522 | 431/38823 | 9.88E-21 | 1.15E-18 | 9.69E-19 | 121 | BP | Ba5B | Metabolism |
| GO:0009123 | nucleoside monophosphate metabolic process | 7/4522 | 18/38823 | 2.86E-03 | 2.96E-02 | 2.49E-02 | 7 | BP | Ba5B | Metabolism |
| GO:0009126 | purine nucleoside monophosphate metabolic process | 6/4522 | 11/38823 | 6.82E-04 | 1.02E-02 | 8.55E-03 | 6 | BP | Ba5B | Metabolism |
| GO:0009141 | nucleoside triphosphate metabolic process | 8/4522 | 15/38823 | 1.01E-04 | 2.06E-03 | 1.73E-03 | 8 | BP | Ba5B | Metabolism |

| | | | | | | | | | | |
|---|---|---|---|---|---|---|---|---|---|---|
| GO:0009144 | purine nucleoside triphosphate metabolic process | 7/4522 | 13/38823 | 2.62E-04 | 4.67E-03 | 3.93E-03 | 7 | BP | Ba5B | Metabolism |
| GO:0009150 | purine ribonucleotide metabolic process | 9/4522 | 25/38823 | 1.39E-03 | 1.67E-02 | 1.41E-02 | 9 | BP | Ba5B | Metabolism |
| GO:0009167 | purine ribonucleoside monophosphate metabolic process | 6/4522 | 11/38823 | 6.82E-04 | 1.02E-02 | 8.55E-03 | 6 | BP | Ba5B | Metabolism |
| GO:0009205 | purine ribonucleoside triphosphate metabolic process | 7/4522 | 13/38823 | 2.62E-04 | 4.67E-03 | 3.93E-03 | 7 | BP | Ba5B | Metabolism |
| GO:0009314 | response to radiation | 76/4522 | 321/38823 | 1.07E-09 | 7.11E-08 | 5.98E-08 | 76 | BP | Ba5B | Defense |
| GO:0009416 | response to light stimulus | 74/4522 | 312/38823 | 1.62E-09 | 1.03E-07 | 8.65E-08 | 74 | BP | Ba5B | Defense |
| GO:0009635 | response to herbicide | 7/4522 | 15/38823 | 7.94E-04 | 1.14E-02 | 9.55E-03 | 7 | BP | Ba5B | Defense |
| GO:0009642 | response to light intensity | 50/4522 | 98/38823 | 1.27E-21 | 1.98E-19 | 1.67E-19 | 50 | BP | Ba5B | Defense |
| GO:0009697 | salicylic acid biosynthetic process | 9/4522 | 10/38823 | 3.51E-08 | 1.85E-06 | 1.56E-06 | 9 | BP | Ba5B | Metabolism |
| GO:0009772 | photosynthetic electron transport in photosystem II | 7/4522 | 11/38823 | 6.23E-05 | 1.34E-03 | 1.13E-03 | 7 | BP | Ba5B | Transport |
| GO:0009893 | positive regulation of metabolic process | 47/4522 | 257/38823 | 1.16E-03 | 1.53E-02 | 1.28E-02 | 47 | BP | Ba5B | Metabolism |
| GO:0009955 | adaxial/abaxial pattern specification | 5/4522 | 10/38823 | 3.25E-03 | 3.23E-02 | 2.72E-02 | 5 | BP | Ba5B | Other |
| GO:0010023 | proanthocyanidin biosynthetic process | 6/4522 | 15/38823 | 4.89E-03 | 4.58E-02 | 3.85E-02 | 6 | BP | Ba5B | Metabolism |
| GO:0010109 | regulation of photosynthesis | 9/4522 | 28/38823 | 3.40E-03 | 3.31E-02 | 2.79E-02 | 9 | BP | Ba5B | Regulation |
| GO:0010345 | suberin biosynthetic process | 9/4522 | 25/38823 | 1.39E-03 | 1.67E-02 | 1.41E-02 | 9 | BP | Ba5B | Metabolism |
| GO:0010380 | regulation of chlorophyll biosynthetic process | 16/4522 | 39/38823 | 2.99E-06 | 8.73E-05 | 7.34E-05 | 16 | BP | Ba5B | Metabolism |
| GO:0010467 | gene expression | 47/4522 | 216/38823 | 1.66E-05 | 4.30E-04 | 3.61E-04 | 47 | BP | Ba5B | Other |
| GO:0010498 | proteasomal protein catabolic process | 26/4522 | 125/38823 | 2.30E-03 | 2.48E-02 | 2.09E-02 | 26 | BP | Ba5B | Other |
| GO:0015074 | DNA integration | 145/4522 | 418/38823 | 2.13E-35 | 2.98E-32 | 2.51E-32 | 145 | BP | Ba5B | Other |
| GO:0015833 | peptide transport | 24/4522 | 109/38823 | 1.50E-03 | 1.75E-02 | 1.47E-02 | 24 | BP | Ba5B | Transport |
| GO:0015977 | carbon fixation | 10/4522 | 24/38823 | 1.89E-04 | 3.53E-03 | 2.97E-03 | 10 | BP | Ba5B | Other |
| GO:0015994 | chlorophyll metabolic process | 9/4522 | 21/38823 | 3.09E-04 | 5.27E-03 | 4.43E-03 | 9 | BP | Ba5B | Metabolism |
| GO:0016053 | organic acid biosynthetic process | 25/4522 | 122/38823 | 3.40E-03 | 3.31E-02 | 2.79E-02 | 25 | BP | Ba5B | Metabolism |
| GO:0016072 | rRNA metabolic process | 17/4522 | 46/38823 | 8.01E-06 | 2.24E-04 | 1.89E-04 | 17 | BP | Ba5B | Metabolism |
| GO:0016310 | phosphorylation | 53/4522 | 296/38823 | 9.78E-04 | 1.35E-02 | 1.13E-02 | 53 | BP | Ba5B | Other |
| GO:0016999 | antibiotic metabolic process | 17/4522 | 40/38823 | 8.09E-07 | 2.80E-05 | 2.35E-05 | 17 | BP | Ba5B | Metabolism |
| GO:0017000 | antibiotic biosynthetic process | 9/4522 | 10/38823 | 3.51E-08 | 1.85E-06 | 1.56E-06 | 9 | BP | Ba5B | Metabolism |

| | | | | | | | | | | |
|---|---|---|---|---|---|---|---|---|---|---|
| GO:0018958 | phenol-containing compound metabolic process | 17/4522 | 39/38823 | 5.22E-07 | 2.15E-05 | 1.81E-05 | 17 | BP | Ba5B | Metabolism |
| GO:0019216 | regulation of lipid metabolic process | 16/4522 | 39/38823 | 2.99E-06 | 8.73E-05 | 7.34E-05 | 16 | BP | Ba5B | Metabolism |
| GO:0019637 | organophosphate metabolic process | 22/4522 | 84/38823 | 1.87E-04 | 3.51E-03 | 2.95E-03 | 22 | BP | Ba5B | Metabolism |
| GO:0019684 | photosynthesis, light reaction | 18/4522 | 55/38823 | 3.02E-05 | 7.41E-04 | 6.23E-04 | 18 | BP | Ba5B | Other |
| GO:0019725 | cellular homeostasis | 21/4522 | 95/38823 | 2.72E-03 | 2.83E-02 | 2.38E-02 | 21 | BP | Ba5B | Other |
| GO:0019747 | regulation of isoprenoid metabolic process | 13/4522 | 30/38823 | 1.24E-05 | 3.29E-04 | 2.76E-04 | 13 | BP | Ba5B | Metabolism |
| GO:0019752 | carboxylic acid metabolic process | 46/4522 | 249/38823 | 1.05E-03 | 1.41E-02 | 1.19E-02 | 46 | BP | Ba5B | Metabolism |
| GO:0019915 | lipid storage | 13/4522 | 34/38823 | 6.14E-05 | 1.34E-03 | 1.13E-03 | 13 | BP | Ba5B | Other |
| GO:0022406 | membrane docking | 6/4522 | 13/38823 | 2.06E-03 | 2.24E-02 | 1.89E-02 | 6 | BP | Ba5B | Other |
| GO:0022607 | cellular component assembly | 37/4522 | 183/38823 | 5.67E-04 | 8.97E-03 | 7.54E-03 | 37 | BP | Ba5B | Other |
| GO:0022613 | ribonucleoprotein complex | 24/4522 | 71/38823 | 7.84E-07 | 2.80E-05 | 2.35E-05 | 24 | BP | Ba5B | Other |
| GO:0022618 | protein-RNA complex assembly | 9/4522 | 22/38823 | 4.68E-04 | 7.50E-03 | 6.30E-03 | 9 | BP | Ba5B | Other |
| GO:0022900 | electron transport chain | 13/4522 | 46/38823 | 1.76E-03 | 2.01E-02 | 1.69E-02 | 13 | BP | Ba5B | Transport |
| GO:0030003 | intracellular monoatomic cation homeostasis | 15/4522 | 57/38823 | 1.78E-03 | 2.02E-02 | 1.70E-02 | 15 | BP | Ba5B | Other |
| GO:0030641 | regulation of cellular pH | 12/4522 | 30/38823 | 7.00E-05 | 1.49E-03 | 1.25E-03 | 12 | BP | Ba5B | Regulation |
| GO:0030656 | regulation of vitamin metabolic process | 6/4522 | 11/38823 | 6.82E-04 | 1.02E-02 | 8.55E-03 | 6 | BP | Ba5B | Metabolism |
| GO:0030951 | establishment or maintenance of microtubule cytoskeleton polarity | 7/4522 | 15/38823 | 7.94E-04 | 1.14E-02 | 9.55E-03 | 7 | BP | Ba5B | Other |
| GO:0031400 | negative regulation of protein modification process | 6/4522 | 14/38823 | 3.25E-03 | 3.23E-02 | 2.72E-02 | 6 | BP | Ba5B | Regulation |
| GO:0031537 | regulation of anthocyanin metabolic process | 7/4522 | 17/38823 | 1.94E-03 | 2.14E-02 | 1.80E-02 | 7 | BP | Ba5B | Metabolism |
| GO:0031540 | regulation of anthocyanin biosynthetic process | 7/4522 | 16/38823 | 1.27E-03 | 1.60E-02 | 1.34E-02 | 7 | BP | Ba5B | Metabolism |
| GO:0031667 | response to nutrient levels | 19/4522 | 76/38823 | 9.25E-04 | 1.29E-02 | 1.08E-02 | 19 | BP | Ba5B | Defense |
| GO:0032101 | regulation of response to external stimulus | 13/4522 | 40/38823 | 4.06E-04 | 6.77E-03 | 5.70E-03 | 13 | BP | Ba5B | Defense |
| GO:0032196 | transposition | 88/4522 | 234/38823 | 7.92E-25 | 3.70E-22 | 3.11E-22 | 88 | BP | Ba5B | Other |
| GO:0032197 | retrotransposition | 88/4522 | 234/38823 | 7.92E-25 | 3.70E-22 | 3.11E-22 | 88 | BP | Ba5B | Other |
| GO:0032264 | IMP salvage | 9/4522 | 23/38823 | 6.90E-04 | 1.02E-02 | 8.61E-03 | 9 | BP | Ba5B | Other |
| GO:0032392 | DNA geometric change | 7/4522 | 14/38823 | 4.71E-04 | 7.50E-03 | 6.30E-03 | 7 | BP | Ba5B | Other |

| | | | | | | | | | | |
|---|---|---|---|---|---|---|---|---|---|---|
| GO:0032787 | monocarboxylic acid metabolic process | 25/4522 | 98/38823 | 1.12E-04 | 2.22E-03 | 1.86E-03 | 25 | BP | Ba5B | Metabolism |
| GO:0033013 | tetrapyrrole metabolic process | 10/4522 | 20/38823 | 2.77E-05 | 6.93E-04 | 5.83E-04 | 10 | BP | Ba5B | Metabolism |
| GO:0033015 | tetrapyrrole catabolic process | 8/4522 | 17/38823 | 3.09E-04 | 5.27E-03 | 4.43E-03 | 8 | BP | Ba5B | Other |
| GO:0033036 | macromolecule localization | 38/4522 | 193/38823 | 8.19E-04 | 1.16E-02 | 9.74E-03 | 38 | BP | Ba5B | Other |
| GO:0034220 | monoatomic ion transmembrane transport | 40/4522 | 212/38823 | 1.43E-03 | 1.71E-02 | 1.44E-02 | 40 | BP | Ba5B | Transport |
| GO:0034389 | lipid droplet organization | 13/4522 | 27/38823 | 2.92E-06 | 8.73E-05 | 7.34E-05 | 13 | BP | Ba5B | Other |
| GO:0034654 | nucleobase-containing compound biosynthetic process | 98/4522 | 321/38823 | 8.12E-20 | 8.12E-18 | 6.83E-18 | 98 | BP | Ba5B | Metabolism |
| GO:0034655 | nucleobase-containing compound catabolic process | 9/4522 | 19/38823 | 1.21E-04 | 2.36E-03 | 1.99E-03 | 9 | BP | Ba5B | Other |
| GO:0035336 | long-chain fatty-acyl-CoA metabolic process | 9/4522 | 25/38823 | 1.39E-03 | 1.67E-02 | 1.41E-02 | 9 | BP | Ba5B | Metabolism |
| GO:0035556 | intracellular signal transduction | 30/4522 | 154/38823 | 3.21E-03 | 3.23E-02 | 2.72E-02 | 30 | BP | Ba5B | Other |
| GO:0035601 | protein deacylation | 6/4522 | 14/38823 | 3.25E-03 | 3.23E-02 | 2.72E-02 | 6 | BP | Ba5B | Other |
| GO:0042254 | ribosome biogenesis | 23/4522 | 100/38823 | 9.99E-04 | 1.36E-02 | 1.14E-02 | 23 | BP | Ba5B | Other |
| GO:0042537 | benzene-containing compound metabolic process | 11/4522 | 24/38823 | 3.08E-05 | 7.44E-04 | 6.26E-04 | 11 | BP | Ba5B | Metabolism |
| GO:0042592 | homeostatic process | 36/4522 | 196/38823 | 3.79E-03 | 3.66E-02 | 3.08E-02 | 36 | BP | Ba5B | Other |
| GO:0042886 | amide transport | 25/4522 | 112/38823 | 9.82E-04 | 1.35E-02 | 1.13E-02 | 25 | BP | Ba5B | Transport |
| GO:0043043 | peptide biosynthetic process | 15/4522 | 58/38823 | 2.15E-03 | 2.32E-02 | 1.95E-02 | 15 | BP | Ba5B | Metabolism |
| GO:0043436 | oxoacid metabolic process | 43/4522 | 220/38823 | 4.58E-04 | 7.45E-03 | 6.27E-03 | 43 | BP | Ba5B | Metabolism |
| GO:0043603 | amide metabolic process | 19/4522 | 76/38823 | 9.25E-04 | 1.29E-02 | 1.08E-02 | 19 | BP | Ba5B | Metabolism |
| GO:0043604 | amide biosynthetic process | 18/4522 | 65/38823 | 3.32E-04 | 5.61E-03 | 4.71E-03 | 18 | BP | Ba5B | Metabolism |
| GO:0044085 | cellular component biogenesis | 57/4522 | 264/38823 | 2.90E-06 | 8.73E-05 | 7.34E-05 | 57 | BP | Ba5B | Other |
| GO:0044281 | small molecule metabolic process | 60/4522 | 324/38823 | 1.92E-04 | 3.55E-03 | 2.99E-03 | 60 | BP | Ba5B | Metabolism |
| GO:0044283 | small molecule biosynthetic process | 32/4522 | 172/38823 | 4.99E-03 | 4.64E-02 | 3.90E-02 | 32 | BP | Ba5B | Metabolism |
| GO:0045184 | establishment of protein localization | 24/4522 | 106/38823 | 9.94E-04 | 1.36E-02 | 1.14E-02 | 24 | BP | Ba5B | Other |
| GO:0045828 | positive regulation of isoprenoid metabolic process | 13/4522 | 30/38823 | 1.24E-05 | 3.29E-04 | 2.76E-04 | 13 | BP | Ba5B | Metabolism |
| GO:0045834 | positive regulation of lipid metabolic process | 13/4522 | 30/38823 | 1.24E-05 | 3.29E-04 | 2.76E-04 | 13 | BP | Ba5B | Metabolism |
| GO:0045851 | pH reduction | 6/4522 | 15/38823 | 4.89E-03 | 4.58E-02 | 3.85E-02 | 6 | BP | Ba5B | Other |
| GO:0046136 | positive regulation of vitamin metabolic process | 6/4522 | 11/38823 | 6.82E-04 | 1.02E-02 | 8.55E-03 | 6 | BP | Ba5B | Metabolism |

| | | | | | | | | | | |
|---|---|---|---|---|---|---|---|---|---|---|
| GO:0046149 | pigment catabolic process | 8/4522 | 16/38823 | 1.82E-04 | 3.44E-03 | 2.90E-03 | 8 | BP | Ba5B | Other |
| GO:0046189 | phenol-containing compound biosynthetic process | 15/4522 | 29/38823 | 1.50E-07 | 7.37E-06 | 6.19E-06 | 15 | BP | Ba5B | Metabolism |
| GO:0046394 | carboxylic acid biosynthetic process | 25/4522 | 122/38823 | 3.40E-03 | 3.31E-02 | 2.79E-02 | 25 | BP | Ba5B | Metabolism |
| GO:0046889 | positive regulation of lipid biosynthetic process | 13/4522 | 30/38823 | 1.24E-05 | 3.29E-04 | 2.76E-04 | 13 | BP | Ba5B | Metabolism |
| GO:0046890 | regulation of lipid biosynthetic process | 13/4522 | 32/38823 | 2.88E-05 | 7.13E-04 | 6.00E-04 | 13 | BP | Ba5B | Metabolism |
| GO:0048878 | chemical homeostasis | 22/4522 | 101/38823 | 2.65E-03 | 2.77E-02 | 2.33E-02 | 22 | BP | Ba5B | Other |
| GO:0050790 | regulation of catalytic activity | 22/4522 | 100/38823 | 2.33E-03 | 2.50E-02 | 2.10E-02 | 22 | BP | Ba5B | Regulation |
| GO:0050801 | monoatomic ion homeostasis | 22/4522 | 95/38823 | 1.16E-03 | 1.53E-02 | 1.28E-02 | 22 | BP | Ba5B | Other |
| GO:0051017 | actin filament bundle assembly | 12/4522 | 39/38823 | 1.16E-03 | 1.53E-02 | 1.28E-02 | 12 | BP | Ba5B | Other |
| GO:0051452 | intracellular pH reduction | 6/4522 | 15/38823 | 4.89E-03 | 4.58E-02 | 3.85E-02 | 6 | BP | Ba5B | Other |
| GO:0051453 | regulation of intracellular pH | 14/4522 | 54/38823 | 2.91E-03 | 2.99E-02 | 2.52E-02 | 14 | BP | Ba5B | Regulation |
| GO:0051639 | actin filament network formation | 10/4522 | 29/38823 | 1.12E-03 | 1.49E-02 | 1.25E-02 | 10 | BP | Ba5B | Other |
| GO:0051641 | cellular localization | 35/4522 | 168/38823 | 4.43E-04 | 7.31E-03 | 6.15E-03 | 35 | BP | Ba5B | Other |
| GO:0055080 | monoatomic cation homeostasis | 17/4522 | 61/38823 | 4.44E-04 | 7.31E-03 | 6.15E-03 | 17 | BP | Ba5B | Other |
| GO:0055082 | intracellular chemical homeostasis | 15/4522 | 55/38823 | 1.20E-03 | 1.57E-02 | 1.32E-02 | 15 | BP | Ba5B | Other |
| GO:0061024 | membrane organization | 28/4522 | 103/38823 | 1.26E-05 | 3.30E-04 | 2.78E-04 | 28 | BP | Ba5B | Other |
| GO:0061025 | membrane fusion | 9/4522 | 26/38823 | 1.91E-03 | 2.14E-02 | 1.80E-02 | 9 | BP | Ba5B | Other |
| GO:0062013 | positive regulation of small molecule metabolic process | 6/4522 | 15/38823 | 4.89E-03 | 4.58E-02 | 3.85E-02 | 6 | BP | Ba5B | Metabolism |
| GO:0065009 | regulation of molecular function | 24/4522 | 115/38823 | 3.17E-03 | 3.23E-02 | 2.72E-02 | 24 | BP | Ba5B | Regulation |
| GO:0070925 | organelle assembly | 12/4522 | 40/38823 | 1.49E-03 | 1.75E-02 | 1.47E-02 | 12 | BP | Ba5B | Other |
| GO:0071826 | protein-RNA complex organization | 9/4522 | 20/38823 | 1.97E-04 | 3.62E-03 | 3.05E-03 | 9 | BP | Ba5B | Other |
| GO:0071897 | DNA biosynthetic process | 89/4522 | 247/38823 | 1.41E-23 | 4.39E-21 | 3.69E-21 | 89 | BP | Ba5B | Metabolism |
| GO:0072330 | monocarboxylic acid biosynthetic process | 15/4522 | 52/38823 | 6.35E-04 | 9.94E-03 | 8.36E-03 | 15 | BP | Ba5B | Metabolism |
| GO:0080177 | plastoglobule organization | 7/4522 | 16/38823 | 1.27E-03 | 1.60E-02 | 1.34E-02 | 7 | BP | Ba5B | Other |
| GO:0080183 | response to photooxidative stress | 7/4522 | 18/38823 | 2.86E-03 | 2.96E-02 | 2.49E-02 | 7 | BP | Ba5B | Defense |
| GO:0090056 | regulation of chlorophyll metabolic process | 16/4522 | 39/38823 | 2.99E-06 | 8.73E-05 | 7.34E-05 | 16 | BP | Ba5B | Metabolism |
| GO:0090174 | organelle membrane fusion | 7/4522 | 16/38823 | 1.27E-03 | 1.60E-02 | 1.34E-02 | 7 | BP | Ba5B | Other |
| GO:0098542 | defense response to other organism | 56/4522 | 325/38823 | 1.79E-03 | 2.03E-02 | 1.70E-02 | 56 | BP | Ba5B | Defense |
| GO:0098655 | monoatomic cation transmembrane transport | 35/4522 | 132/38823 | 2.08E-06 | 6.40E-05 | 5.39E-05 | 35 | BP | Ba5B | Transport |

| | | | | | | | | | | |
|---|---|---|---|---|---|---|---|---|---|---|
| GO:0098660 | inorganic ion transmembrane | 33/4522 | 112/38823 | 3.00E-07 | 1.33E-05 | 1.12E-05 | 33 | BP | Ba5B | Transport |
| GO:0098662 | inorganic cation transmembrane transport | 65/4522 | 196/38823 | 1.43E-15 | 1.15E-13 | 9.64E-14 | 65 | BP | Ba5B | Transport |
| GO:0098732 | macromolecule deacylation | 6/4522 | 14/38823 | 3.25E-03 | 3.23E-02 | 2.72E-02 | 6 | BP | Ba5B | Other |
| GO:0098771 | inorganic ion homeostasis | 17/4522 | 63/38823 | 6.67E-04 | 1.02E-02 | 8.55E-03 | 17 | BP | Ba5B | Other |
| GO:0140056 | organelle localization by membrane tethering | 6/4522 | 13/38823 | 2.06E-03 | 2.24E-02 | 1.89E-02 | 6 | BP | Ba5B | Other |
| GO:1900150 | regulation of defense response to fungus | 11/4522 | 26/38823 | 7.63E-05 | 1.58E-03 | 1.33E-03 | 11 | BP | Ba5B | Defense |
| GO:1900424 | regulation of defense response to bacterium | 9/4522 | 14/38823 | 4.52E-06 | 1.29E-04 | 1.09E-04 | 9 | BP | Ba5B | Defense |
| GO:1901401 | regulation of tetrapyrrole metabolic process | 17/4522 | 40/38823 | 8.09E-07 | 2.80E-05 | 2.35E-05 | 17 | BP | Ba5B | Metabolism |
| GO:1901403 | positive regulation of tetrapyrrole metabolic process | 14/4522 | 27/38823 | 3.77E-07 | 1.60E-05 | 1.35E-05 | 14 | BP | Ba5B | Metabolism |
| GO:1901463 | regulation of tetrapyrrole biosynthetic process | 17/4522 | 40/38823 | 8.09E-07 | 2.80E-05 | 2.35E-05 | 17 | BP | Ba5B | Metabolism |
| GO:1901465 | positive regulation of tetrapyrrole biosynthetic process | 14/4522 | 27/38823 | 3.77E-07 | 1.60E-05 | 1.35E-05 | 14 | BP | Ba5B | Metabolism |
| GO:1901562 | response to paraquat | 7/4522 | 15/38823 | 7.94E-04 | 1.14E-02 | 9.55E-03 | 7 | BP | Ba5B | Defense |
| GO:1902171 | regulation of tocopherol cyclase activity | 7/4522 | 16/38823 | 1.27E-03 | 1.60E-02 | 1.34E-02 | 7 | BP | Ba5B | Regulation |
| GO:1902326 | positive regulation of chlorophyll biosynthetic process | 13/4522 | 26/38823 | 1.70E-06 | 5.28E-05 | 4.44E-05 | 13 | BP | Ba5B | Metabolism |
| GO:1902600 | proton transmembrane transport | 66/4522 | 258/38823 | 4.44E-10 | 3.03E-08 | 2.55E-08 | 66 | BP | Ba5B | Transport |
| GO:1902930 | regulation of alcohol biosynthetic process | 6/4522 | 14/38823 | 3.25E-03 | 3.23E-02 | 2.72E-02 | 6 | BP | Ba5B | Metabolism |
| GO:1902932 | positive regulation of alcohol biosynthetic process | 6/4522 | 14/38823 | 3.25E-03 | 3.23E-02 | 2.72E-02 | 6 | BP | Ba5B | Metabolism |
| GO:1904143 | positive regulation of carotenoid biosynthetic process | 7/4522 | 15/38823 | 7.94E-04 | 1.14E-02 | 9.55E-03 | 7 | BP | Ba5B | Metabolism |
| GO:1904963 | regulation of phytol biosynthetic process | 6/4522 | 11/38823 | 6.82E-04 | 1.02E-02 | 8.55E-03 | 6 | BP | Ba5B | Metabolism |
| GO:1904964 | positive regulation of phytol biosynthetic process | 6/4522 | 11/38823 | 6.82E-04 | 1.02E-02 | 8.55E-03 | 6 | BP | Ba5B | Metabolism |

| | | | | | | | | | | |
|---|---|---|---|---|---|---|---|---|---|---|
| GO:1904965 | regulation of vitamin E biosynthetic process | 6/4522 | 11/38823 | 6.82E-04 | 1.02E-02 | 8.55E-03 | 6 | BP | Ba5B | Metabolism |
| GO:1904966 | positive regulation of vitamin E biosynthetic process | 6/4522 | 11/38823 | 6.82E-04 | 1.02E-02 | 8.55E-03 | 6 | BP | Ba5B | Metabolism |
| GO:2000694 | regulation of phragmoplast microtubule organization | 5/4522 | 11/38823 | 5.39E-03 | 4.95E-02 | 4.16E-02 | 5 | BP | Ba5B | Regulation |

Table S10  Summary of RNA-seq read mapping to the reference genome across different infection stages.

| Days post infection | Biological replicate | Sample | Total reads | Mapped reads | Mapping rate | Depth (×) | Genome coverage at ≥ 1× |
|---|---|---|---|---|---|---|---|
| d3 | r1 | A3 | 162,585,985 | 130,988,515 | 80.57% | 183.26 | 12.27% |
| d3 | r2 | B3 | 205,404,902 | 167,946,236 | 81.76% | 192.28 | 13.72% |
| d3 | r3 | D3 | 159,287,586 | 128,732,011 | 80.82% | 220.90 | 11.92% |
| d4 | r1 | A4 | 193,347,110 | 157,706,740 | 81.57% | 272.48 | 12.71% |
| d4 | r2 | B4 | 176,769,601 | 146,774,175 | 83.03% | 169.32 | 12.82% |
| d4 | r3 | D4 | 201,338,086 | 165,854,245 | 82.38% | 270.29 | 14.31% |
| d7 | r1 | A7 | 184,669,624 | 154,723,856 | 83.78% | 199.20 | 14.35% |
| d7 | r2 | B7 | 212,492,282 | 175,245,200 | 82.47% | 211.66 | 16.72% |
| d7 | r3 | D7 | 187,165,813 | 157,094,483 | 83.93% | 199.34 | 16.86% |
| d11 | r1 | A11 | 213,205,612 | 183,120,325 | 85.89% | 184.94 | 16.55% |
| d11 | r2 | B11 | 221,990,967 | 176,712,090 | 79.60% | 152.02 | 14.82% |
| d11 | r3 | D11 | 186,893,658 | 158,523,316 | 84.82% | 154.14 | 16.06% |
| d0 | r1 | CK1 | 184,204,100 | 153,602,485 | 83.39% | 198.04 | 11.73% |
| d0 | r2 | CK2 | 189,397,710 | 156,547,249 | 82.66% | 208.16 | 13.24% |
| d0 | r3 | CK3 | 232,504,550 | 187,540,719 | 80.66% | 255.45 | 12.15% |

Table S11 GO biological process enrichment results for upregulated genes in barberry during *Pst* infection.

| ID | Description | GeneRatio | BgRatio | pvalue | p.adjust | qvalue | Count | Ontology | cluster |
|---|---|---|---|---|---|---|---|---|---|
| GO:0000949 | aromatic amino acid family catabolic process to alcohol via Ehrlich pathway | 8/2455 | 23/38879 | 5.19E-05 | 7.84E-03 | 7.51E-03 | 8 | BP | d3up |
| GO:0006529 | asparagine biosynthetic process | 8/2455 | 15/38879 | 1.08E-06 | 5.50E-04 | 5.27E-04 | 8 | BP | d3up |
| GO:0006571 | tyrosine biosynthetic process | 7/2455 | 17/38879 | 4.39E-05 | 7.84E-03 | 7.51E-03 | 7 | BP | d3up |
| GO:0006865 | amino acid transport | 15/2455 | 91/38879 | 5.56E-04 | 4.79E-02 | 4.59E-02 | 15 | BP | d3up |
| GO:0009738 | abscisic acid-activated signaling pathway | 15/2455 | 56/38879 | 1.33E-06 | 5.50E-04 | 5.27E-04 | 15 | BP | d3up |
| GO:0009873 | ethylene-activated signaling pathway | 13/2455 | 69/38879 | 3.39E-04 | 3.34E-02 | 3.20E-02 | 13 | BP | d3up |
| GO:0031146 | SCF-dependent proteasomal ubiquitin-dependent protein catabolic process | 14/2455 | 66/38879 | 5.30E-05 | 7.84E-03 | 7.51E-03 | 14 | BP | d3up |
| GO:0000949 | aromatic amino acid family catabolic process to alcohol via Ehrlich pathway | 7/1450 | 23/38879 | 1.43E-05 | 1.31E-03 | 1.25E-03 | 7 | BP | d4up |
| GO:0006096 | glycolytic process | 12/1450 | 109/38879 | 7.74E-04 | 2.95E-02 | 2.80E-02 | 12 | BP | d4up |
| GO:0006571 | tyrosine biosynthetic process | 8/1450 | 17/38879 | 6.61E-08 | 1.72E-05 | 1.64E-05 | 8 | BP | d4up |
| GO:0006749 | glutathione metabolic process | 15/1450 | 80/38879 | 2.38E-07 | 5.43E-05 | 5.17E-05 | 15 | BP | d4up |
| GO:0007005 | mitochondrion organization | 13/1450 | 83/38879 | 1.19E-05 | 1.21E-03 | 1.15E-03 | 13 | BP | d4up |
| GO:0008299 | isoprenoid biosynthetic process | 14/1450 | 125/38879 | 2.45E-04 | 1.31E-02 | 1.25E-02 | 14 | BP | d4up |
| GO:0009094 | L-phenylalanine biosynthetic process | 4/1450 | 11/38879 | 5.15E-04 | 2.30E-02 | 2.18E-02 | 4 | BP | d4up |
| GO:0009738 | abscisic acid-activated signaling pathway | 11/1450 | 56/38879 | 5.94E-06 | 6.39E-04 | 6.08E-04 | 11 | BP | d4up |
| GO:0010200 | response to chitin | 6/1450 | 29/38879 | 6.05E-04 | 2.63E-02 | 2.50E-02 | 6 | BP | d4up |
| GO:0015936 | coenzyme A metabolic process | 9/1450 | 52/38879 | 1.18E-04 | 7.97E-03 | 7.59E-03 | 9 | BP | d4up |
| GO:0016126 | sterol biosynthetic process | 13/1450 | 77/38879 | 5.10E-06 | 6.21E-04 | 5.91E-04 | 13 | BP | d4up |
| GO:0030198 | extracellular matrix organization | 8/1450 | 33/38879 | 2.22E-05 | 1.93E-03 | 1.83E-03 | 8 | BP | d4up |
| GO:0030574 | collagen catabolic process | 8/1450 | 22/38879 | 7.36E-07 | 1.37E-04 | 1.30E-04 | 8 | BP | d4up |
| GO:0042026 | protein refolding | 7/1450 | 41/38879 | 7.31E-04 | 2.91E-02 | 2.77E-02 | 7 | BP | d4up |
| GO:0051123 | RNA polymerase II preinitiation complex assembly | 6/1450 | 25/38879 | 2.56E-04 | 1.34E-02 | 1.27E-02 | 6 | BP | d4up |

| ID | Description | GeneRatio | BgRatio | pvalue | p.adjust | qvalue | Count | Ontology | cluster |
|---|---|---|---|---|---|---|---|---|---|
| GO:0098542 | defense response to other organism | 27/1450 | 313/38879 | 5.39E-05 | 4.28E-03 | 4.07E-03 | 27 | BP | d4up |
| GO:0006032 | chitin catabolic process | 8/2146 | 39/38879 | 1.12E-03 | 4.26E-02 | 3.97E-02 | 8 | BP | d7up |
| GO:0006571 | tyrosine biosynthetic process | 7/2146 | 17/38879 | 1.84E-05 | 1.74E-03 | 1.62E-03 | 7 | BP | d7up |
| GO:0006749 | glutathione metabolic process | 19/2146 | 80/38879 | 5.23E-08 | 1.65E-05 | 1.54E-05 | 19 | BP | d7up |
| GO:0009073 | aromatic amino acid family biosynthetic process | 9/2146 | 46/38879 | 8.04E-04 | 3.71E-02 | 3.46E-02 | 9 | BP | d7up |
| GO:0009410 | response to xenobiotic stimulus | 31/2146 | 243/38879 | 1.31E-05 | 1.38E-03 | 1.28E-03 | 31 | BP | d7up |
| GO:0009611 | response to wounding | 14/2146 | 96/38879 | 7.89E-04 | 3.71E-02 | 3.46E-02 | 14 | BP | d7up |
| GO:0009738 | abscisic acid-activated signaling pathway | 11/2146 | 56/38879 | 2.10E-04 | 1.33E-02 | 1.24E-02 | 11 | BP | d7up |
| GO:0009873 | ethylene-activated signaling pathway | 15/2146 | 69/38879 | 4.28E-06 | 5.41E-04 | 5.04E-04 | 15 | BP | d7up |
| GO:0010112 | regulation of systemic acquired resistance | 8/2146 | 14/38879 | 1.89E-07 | 3.59E-05 | 3.34E-05 | 8 | BP | d7up |
| GO:0010200 | response to chitin | 7/2146 | 29/38879 | 8.23E-04 | 3.71E-02 | 3.46E-02 | 7 | BP | d7up |
| GO:0015936 | coenzyme A metabolic process | 11/2146 | 52/38879 | 1.04E-04 | 7.07E-03 | 6.59E-03 | 11 | BP | d7up |
| GO:0016126 | sterol biosynthetic process | 12/2146 | 77/38879 | 1.01E-03 | 4.04E-02 | 3.76E-02 | 12 | BP | d7up |
| GO:0016998 | cell wall macromolecule catabolic process | 7/2146 | 30/38879 | 1.02E-03 | 4.04E-02 | 3.76E-02 | 7 | BP | d7up |
| GO:0030198 | extracellular matrix organization | 10/2146 | 33/38879 | 7.37E-06 | 8.73E-04 | 8.14E-04 | 10 | BP | d7up |
| GO:0030574 | collagen catabolic process | 10/2146 | 22/38879 | 9.01E-08 | 2.44E-05 | 2.27E-05 | 10 | BP | d7up |
| GO:0045087 | innate immune response | 20/2146 | 164/38879 | 7.50E-04 | 3.71E-02 | 3.46E-02 | 20 | BP | d7up |
| GO:0048544 | recognition of pollen | 12/2146 | 71/38879 | 4.77E-04 | 2.44E-02 | 2.28E-02 | 12 | BP | d7up |
| GO:0050832 | defense response to fungus | 17/2146 | 119/38879 | 2.93E-04 | 1.67E-02 | 1.55E-02 | 17 | BP | d7up |
| GO:0071483 | cellular response to blue light | 5/2146 | 15/38879 | 9.60E-04 | 3.96E-02 | 3.69E-02 | 5 | BP | d7up |
| GO:0080142 | regulation of salicylic acid biosynthetic process | 7/2146 | 18/38879 | 2.87E-05 | 2.36E-03 | 2.20E-03 | 7 | BP | d7up |
| GO:0098542 | defense response to other organism | 40/2146 | 313/38879 | 7.56E-07 | 1.30E-04 | 1.21E-04 | 40 | BP | d7up |
| GO:0000949 | aromatic amino acid family catabolic process to alcohol via Ehrlich pathway | 9/4119 | 23/38879 | 3.40E-04 | 1.21E-02 | 1.13E-02 | 9 | BP | d11up |
| GO:0002229 | defense response to oomycetes | 7/4119 | 14/38879 | 2.60E-04 | 9.72E-03 | 9.03E-03 | 7 | BP | d11up |
| GO:0002239 | response to oomycetes | 9/4119 | 28/38879 | 1.76E-03 | 4.34E-02 | 4.03E-02 | 9 | BP | d11up |

| ID | Description | GeneRatio | BgRatio | pvalue | p.adjust | qvalue | Count | Ontology | cluster |
|---|---|---|---|---|---|---|---|---|---|
| GO:0006002 | fructose 6-phosphate metabolic process | 13/4119 | 27/38879 | 9.90E-07 | 2.15E-04 | 2.00E-04 | 13 | BP | d11up |
| GO:0006096 | glycolytic process | 43/4119 | 109/38879 | 3.45E-15 | 2.06E-12 | 1.92E-12 | 43 | BP | d11up |
| GO:0006470 | protein dephosphorylation | 38/4119 | 213/38879 | 9.48E-04 | 2.70E-02 | 2.51E-02 | 38 | BP | d11up |
| GO:0006486 | protein glycosylation | 35/4119 | 169/38879 | 8.03E-05 | 5.64E-03 | 5.24E-03 | 35 | BP | d11up |
| GO:0006538 | L-glutamate catabolic process | 6/4119 | 12/38879 | 7.39E-04 | 2.18E-02 | 2.03E-02 | 6 | BP | d11up |
| GO:0006635 | fatty acid beta-oxidation | 20/4119 | 77/38879 | 1.13E-04 | 6.92E-03 | 6.43E-03 | 20 | BP | d11up |
| GO:0006749 | glutathione metabolic process | 19/4119 | 80/38879 | 5.68E-04 | 1.89E-02 | 1.75E-02 | 19 | BP | d11up |
| GO:0007166 | cell surface receptor signaling pathway | 22/4119 | 96/38879 | 3.73E-04 | 1.31E-02 | 1.22E-02 | 22 | BP | d11up |
| GO:0009073 | aromatic amino acid family biosynthetic process | 17/4119 | 46/38879 | 2.19E-06 | 4.03E-04 | 3.74E-04 | 17 | BP | d11up |
| GO:0009299 | mRNA transcription | 6/4119 | 14/38879 | 1.99E-03 | 4.76E-02 | 4.42E-02 | 6 | BP | d11up |
| GO:0009410 | response to xenobiotic stimulus | 46/4119 | 243/38879 | 7.19E-05 | 5.21E-03 | 4.84E-03 | 46 | BP | d11up |
| GO:0009605 | response to external stimulus | 66/4119 | 403/38879 | 2.39E-04 | 9.51E-03 | 8.83E-03 | 66 | BP | d11up |
| GO:0009607 | response to biotic stimulus | 54/4119 | 298/38879 | 6.20E-05 | 4.89E-03 | 4.54E-03 | 54 | BP | d11up |
| GO:0009620 | response to fungus | 36/4119 | 127/38879 | 2.36E-08 | 7.04E-06 | 6.54E-06 | 36 | BP | d11up |
| GO:0009696 | salicylic acid metabolic process | 13/4119 | 37/38879 | 6.34E-05 | 4.89E-03 | 4.54E-03 | 13 | BP | d11up |
| GO:0009749 | response to glucose | 10/4119 | 25/38879 | 1.28E-04 | 7.41E-03 | 6.88E-03 | 10 | BP | d11up |
| GO:0009751 | response to salicylic acid | 24/4119 | 103/38879 | 1.58E-04 | 8.21E-03 | 7.63E-03 | 24 | BP | d11up |
| GO:0010112 | regulation of systemic acquired resistance | 9/4119 | 14/38879 | 2.03E-06 | 4.03E-04 | 3.74E-04 | 9 | BP | d11up |
| GO:0010268 | brassinosteroid homeostasis | 11/4119 | 35/38879 | 6.91E-04 | 2.16E-02 | 2.01E-02 | 11 | BP | d11up |
| GO:0015749 | monosaccharide transmembrane transport | 9/4119 | 28/38879 | 1.76E-03 | 4.34E-02 | 4.03E-02 | 9 | BP | d11up |
| GO:0015977 | carbon fixation | 8/4119 | 15/38879 | 5.10E-05 | 4.69E-03 | 4.36E-03 | 8 | BP | d11up |
| GO:0016125 | sterol metabolic process | 18/4119 | 71/38879 | 3.37E-04 | 1.21E-02 | 1.13E-02 | 18 | BP | d11up |
| GO:0016132 | brassinosteroid biosynthetic process | 11/4119 | 38/38879 | 1.48E-03 | 3.88E-02 | 3.61E-02 | 11 | BP | d11up |
| GO:0016999 | antibiotic metabolic process | 9/4119 | 24/38879 | 4.93E-04 | 1.68E-02 | 1.56E-02 | 9 | BP | d11up |
| GO:0017001 | antibiotic catabolic process | 6/4119 | 13/38879 | 1.25E-03 | 3.35E-02 | 3.12E-02 | 6 | BP | d11up |
| GO:0019336 | phenol-containing compound catabolic process | 5/4119 | 10/38879 | 2.12E-03 | 4.97E-02 | 4.62E-02 | 5 | BP | d11up |
| GO:0030198 | extracellular matrix organization | 10/4119 | 33/38879 | 1.64E-03 | 4.22E-02 | 3.92E-02 | 10 | BP | d11up |

| ID | Description | GeneRatio | BgRatio | pvalue | p.adjust | qvalue | Count | Ontology | cluster |
|---|---|---|---|---|---|---|---|---|---|
| GO:0030574 | collagen catabolic process | 10/4119 | 22/38879 | 3.41E-05 | 3.55E-03 | 3.30E-03 | 10 | BP | d11up |
| GO:0031341 | regulation of cell killing | 6/4119 | 10/38879 | 2.03E-04 | 8.21E-03 | 7.63E-03 | 6 | BP | d11up |
| GO:0031343 | positive regulation of cell killing | 6/4119 | 10/38879 | 2.03E-04 | 8.21E-03 | 7.63E-03 | 6 | BP | d11up |
| GO:0042268 | regulation of cytolysis | 6/4119 | 10/38879 | 2.03E-04 | 8.21E-03 | 7.63E-03 | 6 | BP | d11up |
| GO:0042445 | hormone metabolic process | 19/4119 | 81/38879 | 6.70E-04 | 2.16E-02 | 2.01E-02 | 19 | BP | d11up |
| GO:0042537 | benzene-containing compound metabolic process | 8/4119 | 18/38879 | 2.59E-04 | 9.72E-03 | 9.03E-03 | 8 | BP | d11up |
| GO:0042742 | defense response to bacterium | 19/4119 | 87/38879 | 1.67E-03 | 4.24E-02 | 3.94E-02 | 19 | BP | d11up |
| GO:0043207 | response to external biotic stimulus | 54/4119 | 298/38879 | 6.20E-05 | 4.89E-03 | 4.54E-03 | 54 | BP | d11up |
| GO:0043903 | regulation of biological process involved in symbiotic interaction | 7/4119 | 16/38879 | 7.16E-04 | 2.16E-02 | 2.01E-02 | 7 | BP | d11up |
| GO:0045919 | positive regulation of cytolysis | 6/4119 | 10/38879 | 2.03E-04 | 8.21E-03 | 7.63E-03 | 6 | BP | d11up |
| GO:0046244 | salicylic acid catabolic process | 5/4119 | 10/38879 | 2.12E-03 | 4.97E-02 | 4.62E-02 | 5 | BP | d11up |
| GO:0046677 | response to antibiotic | 27/4119 | 137/38879 | 1.10E-03 | 3.09E-02 | 2.87E-02 | 27 | BP | d11up |
| GO:0048544 | recognition of pollen | 22/4119 | 71/38879 | 2.37E-06 | 4.05E-04 | 3.76E-04 | 22 | BP | d11up |
| GO:0050832 | defense response to fungus | 32/4119 | 119/38879 | 5.14E-07 | 1.23E-04 | 1.14E-04 | 32 | BP | d11up |
| GO:0051707 | response to other organism | 54/4119 | 297/38879 | 5.65E-05 | 4.82E-03 | 4.48E-03 | 54 | BP | d11up |
| GO:0051709 | regulation of killing of cells of another organism | 6/4119 | 10/38879 | 2.03E-04 | 8.21E-03 | 7.63E-03 | 6 | BP | d11up |
| GO:0051712 | positive regulation of killing of cells of another organism | 6/4119 | 10/38879 | 2.03E-04 | 8.21E-03 | 7.63E-03 | 6 | BP | d11up |
| GO:0072488 | ammonium transmembrane transport | 9/4119 | 25/38879 | 6.98E-04 | 2.16E-02 | 2.01E-02 | 9 | BP | d11up |
| GO:0080142 | regulation of salicylic acid biosynthetic process | 9/4119 | 18/38879 | 3.31E-05 | 3.55E-03 | 3.30E-03 | 9 | BP | d11up |
| GO:0098542 | defense response to other organism | 76/4119 | 313/38879 | 3.42E-12 | 1.36E-09 | 1.27E-09 | 76 | BP | d11up |
| GO:2000031 | regulation of salicylic acid mediated signaling pathway | 7/4119 | 16/38879 | 7.16E-04 | 2.16E-02 | 2.01E-02 | 7 | BP | d11up |

Table S12 GO biological process enrichment results for downregulated genes in barberry during *Pst* infection.

| ID | Description | GeneRatio | BgRatio | pvalue | p.adjust | qvalue | Count | Ontology | cluster |
|---|---|---|---|---|---|---|---|---|---|
| GO:0006024 | glycosaminoglycan biosynthetic process | 6/2536 | 12/38879 | 5.02E-05 | 1.97E-03 | 1.85E-03 | 6 | BP | d3down |
| GO:0006457 | protein folding | 45/2536 | 310/38879 | 4.21E-07 | 5.52E-05 | 5.17E-05 | 45 | BP | d3down |
| GO:0006914 | autophagy | 15/2536 | 75/38879 | 8.63E-05 | 3.03E-03 | 2.84E-03 | 15 | BP | d3down |
| GO:0009099 | L-valine biosynthetic process | 5/2536 | 14/38879 | 1.43E-03 | 3.56E-02 | 3.34E-02 | 5 | BP | d3down |
| GO:0009249 | protein lipoylation | 8/2536 | 13/38879 | 3.10E-07 | 4.69E-05 | 4.40E-05 | 8 | BP | d3down |
| GO:0009411 | response to UV | 11/2536 | 45/38879 | 1.14E-04 | 3.80E-03 | 3.56E-03 | 11 | BP | d3down |
| GO:0009642 | response to light intensity | 14/2536 | 65/38879 | 6.31E-05 | 2.43E-03 | 2.28E-03 | 14 | BP | d3down |
| GO:0009644 | response to high light intensity | 12/2536 | 45/38879 | 2.19E-05 | 9.80E-04 | 9.19E-04 | 12 | BP | d3down |
| GO:0009658 | chloroplast organization | 18/2536 | 114/38879 | 4.19E-04 | 1.23E-02 | 1.15E-02 | 18 | BP | d3down |
| GO:0009812 | flavonoid metabolic process | 9/2536 | 23/38879 | 7.44E-06 | 4.06E-04 | 3.81E-04 | 9 | BP | d3down |
| GO:0009813 | flavonoid biosynthetic process | 10/2536 | 26/38879 | 2.76E-06 | 2.22E-04 | 2.08E-04 | 10 | BP | d3down |
| GO:0009834 | plant-type secondary cell wall biogenesis | 8/2536 | 30/38879 | 5.18E-04 | 1.43E-02 | 1.34E-02 | 8 | BP | d3down |
| GO:0010029 | regulation of seed germination | 12/2536 | 33/38879 | 5.64E-07 | 6.16E-05 | 5.77E-05 | 12 | BP | d3down |
| GO:0010030 | positive regulation of seed germination | 10/2536 | 20/38879 | 1.38E-07 | 2.71E-05 | 2.54E-05 | 10 | BP | d3down |
| GO:0010114 | response to red light | 8/2536 | 33/38879 | 1.03E-03 | 2.63E-02 | 2.47E-02 | 8 | BP | d3down |
| GO:0010117 | photoprotection | 7/2536 | 15/38879 | 2.01E-05 | 9.41E-04 | 8.82E-04 | 7 | BP | d3down |
| GO:0010206 | photosystem II repair | 9/2536 | 19/38879 | 1.07E-06 | 1.00E-04 | 9.37E-05 | 9 | BP | d3down |
| GO:0010218 | response to far red light | 7/2536 | 23/38879 | 4.82E-04 | 1.35E-02 | 1.27E-02 | 7 | BP | d3down |
| GO:0010224 | response to UV-B | 12/2536 | 40/38879 | 5.78E-06 | 3.55E-04 | 3.33E-04 | 12 | BP | d3down |
| GO:0010380 | regulation of chlorophyll biosynthetic process | 8/2536 | 21/38879 | 3.05E-05 | 1.28E-03 | 1.20E-03 | 8 | BP | d3down |
| GO:0015718 | monocarboxylic acid transport | 5/2536 | 12/38879 | 6.32E-04 | 1.70E-02 | 1.60E-02 | 5 | BP | d3down |
| GO:0015979 | photosynthesis | 23/2536 | 121/38879 | 3.20E-06 | 2.24E-04 | 2.10E-04 | 23 | BP | d3down |
| GO:0019464 | glycine decarboxylation via glycine cleavage system | 8/2536 | 13/38879 | 3.10E-07 | 4.69E-05 | 4.40E-05 | 8 | BP | d3down |
| GO:0034644 | cellular response to UV | 7/2536 | 14/38879 | 1.14E-05 | 5.73E-04 | 5.37E-04 | 7 | BP | d3down |
| GO:0042594 | response to starvation | 17/2536 | 76/38879 | 6.27E-06 | 3.62E-04 | 3.40E-04 | 17 | BP | d3down |
| GO:0042761 | very long-chain fatty acid biosynthetic process | 7/2536 | 24/38879 | 6.42E-04 | 1.71E-02 | 1.60E-02 | 7 | BP | d3down |

| ID | Description | GeneRatio | BgRatio | pvalue | p.adjust | qvalue | Count | Ontology | cluster |
|---|---|---|---|---|---|---|---|---|---|
| GO:0045454 | cell redox homeostasis | 11/2536 | 61/38879 | 1.80E-03 | 4.36E-02 | 4.09E-02 | 11 | BP | d3down |
| GO:0048544 | recognition of pollen | 12/2536 | 71/38879 | 2.02E-03 | 4.85E-02 | 4.55E-02 | 12 | BP | d3down |
| GO:0048582 | positive regulation of post-embryonic development | 11/2536 | 43/38879 | 7.29E-05 | 2.65E-03 | 2.49E-03 | 11 | BP | d3down |
| GO:0070141 | response to UV-A | 7/2536 | 12/38879 | 2.95E-06 | 2.22E-04 | 2.08E-04 | 7 | BP | d3down |
| GO:0071214 | cellular response to abiotic stimulus | 10/2536 | 41/38879 | 2.37E-04 | 7.27E-03 | 6.81E-03 | 10 | BP | d3down |
| GO:0071478 | cellular response to radiation | 8/2536 | 28/38879 | 3.09E-04 | 9.35E-03 | 8.77E-03 | 8 | BP | d3down |
| GO:0071482 | cellular response to light stimulus | 8/2536 | 29/38879 | 4.03E-04 | 1.20E-02 | 1.12E-02 | 8 | BP | d3down |
| GO:0071483 | cellular response to blue light | 7/2536 | 15/38879 | 2.01E-05 | 9.41E-04 | 8.82E-04 | 7 | BP | d3down |
| GO:0071484 | cellular response to light intensity | 7/2536 | 12/38879 | 2.95E-06 | 2.22E-04 | 2.08E-04 | 7 | BP | d3down |
| GO:0071486 | cellular response to high light intensity | 7/2536 | 13/38879 | 6.02E-06 | 3.59E-04 | 3.36E-04 | 7 | BP | d3down |
| GO:0071489 | cellular response to red or far red light | 8/2536 | 18/38879 | 7.84E-06 | 4.17E-04 | 3.91E-04 | 8 | BP | d3down |
| GO:0071490 | cellular response to far red light | 7/2536 | 11/38879 | 1.30E-06 | 1.16E-04 | 1.09E-04 | 7 | BP | d3down |
| GO:0071491 | cellular response to red light | 7/2536 | 10/38879 | 5.02E-07 | 6.16E-05 | 5.77E-05 | 7 | BP | d3down |
| GO:0071492 | cellular response to UV-A | 7/2536 | 12/38879 | 2.95E-06 | 2.22E-04 | 2.08E-04 | 7 | BP | d3down |
| GO:0080167 | response to karrikin | 13/2536 | 60/38879 | 1.07E-04 | 3.69E-03 | 3.46E-03 | 13 | BP | d3down |
| GO:0090056 | regulation of chlorophyll metabolic process | 8/2536 | 21/38879 | 3.05E-05 | 1.28E-03 | 1.20E-03 | 8 | BP | d3down |
| GO:0098869 | cellular oxidant detoxification | 10/2536 | 44/38879 | 4.38E-04 | 1.25E-02 | 1.18E-02 | 10 | BP | d3down |
| GO:0104004 | cellular response to environmental stimulus | 10/2536 | 41/38879 | 2.37E-04 | 7.27E-03 | 6.81E-03 | 10 | BP | d3down |
| GO:1900140 | regulation of seedling development | 12/2536 | 33/38879 | 5.64E-07 | 6.16E-05 | 5.77E-05 | 12 | BP | d3down |
| GO:1901401 | regulation of tetrapyrrole metabolic process | 8/2536 | 23/38879 | 6.54E-05 | 2.43E-03 | 2.28E-03 | 8 | BP | d3down |
| GO:1901463 | regulation of tetrapyrrole biosynthetic process | 8/2536 | 23/38879 | 6.54E-05 | 2.43E-03 | 2.28E-03 | 8 | BP | d3down |
| GO:0001504 | neurotransmitter uptake | 5/1012 | 15/38879 | 2.86E-05 | 1.00E-03 | 9.19E-04 | 5 | BP | d4down |
| GO:0003333 | amino acid transmembrane transport | 12/1012 | 91/38879 | 4.33E-06 | 2.69E-04 | 2.47E-04 | 12 | BP | d4down |
| GO:0005992 | trehalose biosynthetic process | 5/1012 | 29/38879 | 8.36E-04 | 1.30E-02 | 1.19E-02 | 5 | BP | d4down |
| GO:0006811 | monoatomic ion transport | 17/1012 | 296/38879 | 2.13E-03 | 2.97E-02 | 2.72E-02 | 17 | BP | d4down |
| GO:0006820 | monoatomic anion transport | 14/1012 | 218/38879 | 1.85E-03 | 2.60E-02 | 2.39E-02 | 14 | BP | d4down |
| GO:0006835 | dicarboxylic acid transport | 5/1012 | 19/38879 | 1.02E-04 | 2.52E-03 | 2.31E-03 | 5 | BP | d4down |

| ID | Description | GeneRatio | BgRatio | pvalue | p.adjust | qvalue | Count | Ontology | cluster |
|---|---|---|---|---|---|---|---|---|---|
| GO:0006836 | neurotransmitter transport | 5/1012 | 17/38879 | 5.64E-05 | 1.68E-03 | 1.54E-03 | 5 | BP | d4down |
| GO:0006855 | xenobiotic transmembrane transport | 7/1012 | 44/38879 | 1.31E-04 | 2.72E-03 | 2.49E-03 | 7 | BP | d4down |
| GO:0006865 | amino acid transport | 9/1012 | 91/38879 | 6.16E-04 | 1.00E-02 | 9.20E-03 | 9 | BP | d4down |
| GO:0006868 | glutamine transport | 4/1012 | 11/38879 | 1.30E-04 | 2.72E-03 | 2.49E-03 | 4 | BP | d4down |
| GO:0006914 | autophagy | 20/1012 | 75/38879 | 3.52E-15 | 8.03E-13 | 7.36E-13 | 20 | BP | d4down |
| GO:0008643 | carbohydrate transport | 7/1012 | 55/38879 | 5.42E-04 | 9.15E-03 | 8.39E-03 | 7 | BP | d4down |
| GO:0009765 | photosynthesis, light harvesting | 31/1012 | 53/38879 | 1.29E-35 | 1.77E-32 | 1.62E-32 | 31 | BP | d4down |
| GO:0010206 | photosystem II repair | 5/1012 | 19/38879 | 1.02E-04 | 2.52E-03 | 2.31E-03 | 5 | BP | d4down |
| GO:0010207 | photosystem II assembly | 4/1012 | 22/38879 | 2.30E-03 | 3.17E-02 | 2.91E-02 | 4 | BP | d4down |
| GO:0015711 | organic anion transport | 13/1012 | 135/38879 | 5.48E-05 | 1.68E-03 | 1.54E-03 | 13 | BP | d4down |
| GO:0015718 | monocarboxylic acid transport | 3/1012 | 12/38879 | 3.24E-03 | 4.19E-02 | 3.84E-02 | 3 | BP | d4down |
| GO:0015740 | C4-dicarboxylate transport | 5/1012 | 17/38879 | 5.64E-05 | 1.68E-03 | 1.54E-03 | 5 | BP | d4down |
| GO:0015800 | acidic amino acid transport | 5/1012 | 15/38879 | 2.86E-05 | 1.00E-03 | 9.19E-04 | 5 | BP | d4down |
| GO:0015801 | aromatic amino acid transport | 4/1012 | 12/38879 | 1.91E-04 | 3.79E-03 | 3.47E-03 | 4 | BP | d4down |
| GO:0015804 | neutral amino acid transport | 5/1012 | 31/38879 | 1.15E-03 | 1.72E-02 | 1.58E-02 | 5 | BP | d4down |
| GO:0015807 | L-amino acid transport | 5/1012 | 19/38879 | 1.02E-04 | 2.52E-03 | 2.31E-03 | 5 | BP | d4down |
| GO:0015808 | L-alanine transport | 5/1012 | 15/38879 | 2.86E-05 | 1.00E-03 | 9.19E-04 | 5 | BP | d4down |
| GO:0015810 | aspartate transmembrane transport | 4/1012 | 11/38879 | 1.30E-04 | 2.72E-03 | 2.49E-03 | 4 | BP | d4down |
| GO:0015813 | L-glutamate transmembrane transport | 5/1012 | 15/38879 | 2.86E-05 | 1.00E-03 | 9.19E-04 | 5 | BP | d4down |
| GO:0015824 | proline transport | 4/1012 | 15/38879 | 4.95E-04 | 8.58E-03 | 7.86E-03 | 4 | BP | d4down |
| GO:0015825 | L-serine transport | 4/1012 | 11/38879 | 1.30E-04 | 2.72E-03 | 2.49E-03 | 4 | BP | d4down |
| GO:0015827 | tryptophan transport | 4/1012 | 11/38879 | 1.30E-04 | 2.72E-03 | 2.49E-03 | 4 | BP | d4down |
| GO:0015849 | organic acid transport | 12/1012 | 76/38879 | 6.09E-07 | 4.62E-05 | 4.24E-05 | 12 | BP | d4down |
| GO:0015979 | photosynthesis | 31/1012 | 121/38879 | 3.44E-22 | 2.35E-19 | 2.15E-19 | 31 | BP | d4down |
| GO:0015995 | chlorophyll biosynthetic process | 4/1012 | 14/38879 | 3.71E-04 | 6.83E-03 | 6.26E-03 | 4 | BP | d4down |
| GO:0016123 | xanthophyll biosynthetic process | 5/1012 | 15/38879 | 2.86E-05 | 1.00E-03 | 9.19E-04 | 5 | BP | d4down |
| GO:0032328 | alanine transport | 5/1012 | 15/38879 | 2.86E-05 | 1.00E-03 | 9.19E-04 | 5 | BP | d4down |
| GO:0032329 | serine transport | 4/1012 | 11/38879 | 1.30E-04 | 2.72E-03 | 2.49E-03 | 4 | BP | d4down |
| GO:0034220 | monoatomic ion transmembrane transport | 15/1012 | 207/38879 | 3.74E-04 | 6.83E-03 | 6.26E-03 | 15 | BP | d4down |
| GO:0035524 | proline transmembrane transport | 4/1012 | 15/38879 | 4.95E-04 | 8.58E-03 | 7.86E-03 | 4 | BP | d4down |
| GO:0042594 | response to starvation | 22/1012 | 76/38879 | 2.12E-17 | 9.66E-15 | 8.85E-15 | 22 | BP | d4down |
| GO:0042761 | very long-chain fatty acid biosynthetic process | 4/1012 | 24/38879 | 3.20E-03 | 4.19E-02 | 3.84E-02 | 4 | BP | d4down |

| ID | Description | GeneRatio | BgRatio | pvalue | p.adjust | qvalue | Count | Ontology | cluster |
|---|---|---|---|---|---|---|---|---|---|
| GO:0042908 | xenobiotic transport | 7/1012 | 44/38879 | 1.31E-04 | 2.72E-03 | 2.49E-03 | 7 | BP | d4down |
| GO:0043090 | amino acid import | 5/1012 | 21/38879 | 1.70E-04 | 3.42E-03 | 3.14E-03 | 5 | BP | d4down |
| GO:0046942 | carboxylic acid transport | 12/1012 | 76/38879 | 6.09E-07 | 4.62E-05 | 4.24E-05 | 12 | BP | d4down |
| GO:0051938 | L-glutamate import | 5/1012 | 15/38879 | 2.86E-05 | 1.00E-03 | 9.19E-04 | 5 | BP | d4down |
| GO:0071731 | response to nitric oxide | 3/1012 | 11/38879 | 2.48E-03 | 3.33E-02 | 3.05E-02 | 3 | BP | d4down |
| GO:0071732 | cellular response to nitric oxide | 3/1012 | 11/38879 | 2.48E-03 | 3.33E-02 | 3.05E-02 | 3 | BP | d4down |
| GO:0089718 | amino acid import across plasma membrane | 5/1012 | 15/38879 | 2.86E-05 | 1.00E-03 | 9.19E-04 | 5 | BP | d4down |
| GO:0097366 | response to bronchodilator | 3/1012 | 11/38879 | 2.48E-03 | 3.33E-02 | 3.05E-02 | 3 | BP | d4down |
| GO:0098656 | monoatomic anion transmembrane transport | 13/1012 | 111/38879 | 6.66E-06 | 3.80E-04 | 3.48E-04 | 13 | BP | d4down |
| GO:0098657 | import into cell | 5/1012 | 29/38879 | 8.36E-04 | 1.30E-02 | 1.19E-02 | 5 | BP | d4down |
| GO:0098712 | L-glutamate import across plasma membrane | 5/1012 | 15/38879 | 2.86E-05 | 1.00E-03 | 9.19E-04 | 5 | BP | d4down |
| GO:0098739 | import across plasma membrane | 5/1012 | 18/38879 | 7.64E-05 | 2.22E-03 | 2.04E-03 | 5 | BP | d4down |
| GO:1902475 | L-alpha-amino acid transmembrane transport | 5/1012 | 19/38879 | 1.02E-04 | 2.52E-03 | 2.31E-03 | 5 | BP | d4down |
| GO:1903825 | organic acid transmembrane transport | 12/1012 | 76/38879 | 6.09E-07 | 4.62E-05 | 4.24E-05 | 12 | BP | d4down |
| GO:1905039 | carboxylic acid transmembrane transport | 12/1012 | 76/38879 | 6.09E-07 | 4.62E-05 | 4.24E-05 | 12 | BP | d4down |
| GO:0009765 | photosynthesis, light harvesting | 27/823 | 53/38879 | 2.36E-31 | 2.73E-28 | 2.56E-28 | 27 | BP | d7down |
| GO:0010091 | trichome branching | 4/823 | 13/38879 | 1.22E-04 | 1.77E-02 | 1.66E-02 | 4 | BP | d7down |
| GO:0010345 | suberin biosynthetic process | 7/823 | 27/38879 | 1.14E-06 | 2.63E-04 | 2.47E-04 | 7 | BP | d7down |
| GO:0015979 | photosynthesis | 10/823 | 121/38879 | 2.62E-04 | 3.03E-02 | 2.84E-02 | 10 | BP | d7down |
| GO:0016114 | terpenoid biosynthetic process | 12/823 | 134/38879 | 2.98E-05 | 4.92E-03 | 4.62E-03 | 12 | BP | d7down |
| GO:0035336 | long-chain fatty-acyl-CoA metabolic process | 7/823 | 24/38879 | 4.69E-07 | 1.36E-04 | 1.27E-04 | 7 | BP | d7down |
| GO:0042761 | very long-chain fatty acid biosynthetic process | 6/823 | 24/38879 | 8.58E-06 | 1.65E-03 | 1.55E-03 | 6 | BP | d7down |
| GO:0000373 | Group II intron splicing | 9/3974 | 23/38879 | 2.59E-04 | 1.01E-02 | 9.53E-03 | 9 | BP | d11down |
| GO:0005986 | sucrose biosynthetic process | 6/3974 | 10/38879 | 1.66E-04 | 6.93E-03 | 6.55E-03 | 6 | BP | d11down |
| GO:0006779 | porphyrin-containing compound biosynthetic process | 7/3974 | 17/38879 | 8.92E-04 | 2.72E-02 | 2.57E-02 | 7 | BP | d11down |
| GO:0008610 | lipid biosynthetic process | 32/3974 | 168/38879 | 4.04E-04 | 1.52E-02 | 1.44E-02 | 32 | BP | d11down |

| ID | Description | GeneRatio | BgRatio | pvalue | p.adjust | qvalue | Count | Ontology | cluster |
|---|---|---|---|---|---|---|---|---|---|
| GO:0008643 | carbohydrate transport | 14/3974 | 55/38879 | 1.01E-03 | 2.99E-02 | 2.83E-02 | 14 | BP | d11down |
| GO:0009249 | protein lipoylation | 9/3974 | 13/38879 | 5.88E-07 | 4.28E-05 | 4.05E-05 | 9 | BP | d11down |
| GO:0009642 | response to light intensity | 16/3974 | 65/38879 | 6.76E-04 | 2.12E-02 | 2.01E-02 | 16 | BP | d11down |
| GO:0009657 | plastid organization | 28/3974 | 97/38879 | 2.61E-07 | 2.35E-05 | 2.23E-05 | 28 | BP | d11down |
| GO:0009658 | chloroplast organization | 32/3974 | 114/38879 | 7.76E-08 | 7.30E-06 | 6.90E-06 | 32 | BP | d11down |
| GO:0009765 | photosynthesis, light harvesting | 35/3974 | 53/38879 | 1.88E-22 | 8.48E-20 | 8.02E-20 | 35 | BP | d11down |
| GO:0009834 | plant-type secondary cell wall biogenesis | 10/3974 | 30/38879 | 5.38E-04 | 1.87E-02 | 1.77E-02 | 10 | BP | d11down |
| GO:0010020 | chloroplast fission | 16/3974 | 34/38879 | 4.99E-08 | 4.98E-06 | 4.71E-06 | 16 | BP | d11down |
| GO:0010027 | thylakoid membrane organization | 17/3974 | 56/38879 | 2.75E-05 | 1.44E-03 | 1.36E-03 | 17 | BP | d11down |
| GO:0010206 | photosystem II repair | 11/3974 | 19/38879 | 4.34E-07 | 3.63E-05 | 3.43E-05 | 11 | BP | d11down |
| GO:0010207 | photosystem II assembly | 11/3974 | 22/38879 | 3.02E-06 | 2.01E-04 | 1.90E-04 | 11 | BP | d11down |
| GO:0015718 | monocarboxylic acid transport | 6/3974 | 12/38879 | 6.08E-04 | 1.93E-02 | 1.83E-02 | 6 | BP | d11down |
| GO:0015979 | photosynthesis | 67/3974 | 121/38879 | 8.96E-35 | 1.01E-31 | 9.56E-32 | 67 | BP | d11down |
| GO:0015995 | chlorophyll biosynthetic process | 11/3974 | 14/38879 | 3.41E-09 | 3.85E-07 | 3.64E-07 | 11 | BP | d11down |
| GO:0016554 | cytidine to uridine editing | 8/3974 | 20/38879 | 4.81E-04 | 1.75E-02 | 1.65E-02 | 8 | BP | d11down |
| GO:0019432 | triglyceride biosynthetic process | 16/3974 | 48/38879 | 1.26E-05 | 6.80E-04 | 6.43E-04 | 16 | BP | d11down |
| GO:0019464 | glycine decarboxylation via glycine cleavage system | 9/3974 | 13/38879 | 5.88E-07 | 4.28E-05 | 4.05E-05 | 9 | BP | d11down |
| GO:0030244 | cellulose biosynthetic process | 21/3974 | 96/38879 | 6.01E-04 | 1.93E-02 | 1.83E-02 | 21 | BP | d11down |
| GO:0032544 | plastid translation | 7/3974 | 12/38879 | 5.77E-05 | 2.60E-03 | 2.46E-03 | 7 | BP | d11down |
| GO:0034196 | acylglycerol transport | 6/3974 | 12/38879 | 6.08E-04 | 1.93E-02 | 1.83E-02 | 6 | BP | d11down |
| GO:0042026 | protein refolding | 15/3974 | 41/38879 | 6.42E-06 | 3.63E-04 | 3.43E-04 | 15 | BP | d11down |
| GO:0042594 | response to starvation | 18/3974 | 76/38879 | 5.31E-04 | 1.87E-02 | 1.77E-02 | 18 | BP | d11down |
| GO:0042761 | very long-chain fatty acid biosynthetic process | 10/3974 | 24/38879 | 6.24E-05 | 2.76E-03 | 2.61E-03 | 10 | BP | d11down |
| GO:0045017 | glycerolipid biosynthetic process | 16/3974 | 52/38879 | 3.91E-05 | 1.91E-03 | 1.81E-03 | 16 | BP | d11down |
| GO:0045037 | protein import into chloroplast stroma | 12/3974 | 29/38879 | 1.25E-05 | 6.80E-04 | 6.43E-04 | 12 | BP | d11down |
| GO:0045454 | cell redox homeostasis | 15/3974 | 61/38879 | 9.95E-04 | 2.99E-02 | 2.83E-02 | 15 | BP | d11down |
| GO:0045727 | positive regulation of translation | 9/3974 | 25/38879 | 5.36E-04 | 1.87E-02 | 1.77E-02 | 9 | BP | d11down |
| GO:1901259 | chloroplast rRNA processing | 23/3974 | 32/38879 | 1.74E-16 | 5.62E-14 | 5.32E-14 | 23 | BP | d11down |
| GO:1990052 | ER to chloroplast lipid transport | 6/3974 | 12/38879 | 6.08E-04 | 1.93E-02 | 1.83E-02 | 6 | BP | d11down |
